# The neural basis of defensive behaviour evolution in *Peromyscus* mice

**DOI:** 10.1101/2023.07.04.547734

**Authors:** Felix Baier, Katja Reinhard, Victoria Tong, Julie Murmann, Karl Farrow, Hopi E. Hoekstra

## Abstract

Evading imminent predator threat is critical for survival. Effective defensive strategies can vary, even between closely related species. However, the neural basis of such species-specific behaviours is still poorly understood. Here we find that two sister species of deer mice (genus *Peromyscus*) show different responses to the same looming stimulus: *P. maniculatus,* which occupy densely vegetated habitats, predominantly dart to escape, while the open field specialist, *P. polionotus,* pause their movement. This difference arises from species-specific escape thresholds, is largely context-independent, and can be triggered by both visual and auditory threat stimuli. Using immunohistochemistry and electrophysiological recordings, we find that although visual threat activates the superior colliculus in both species, the role of the dorsal periaqueductal gray (dPAG) in driving behaviour differs. While dPAG activity scales with running speed and involves both excitatory and inhibitory neurons in *P. maniculatus*, the dPAG is largely silent in *P. polionotus,* even when darting is triggered. Moreover, optogenetic activation of excitatory dPAG neurons reliably elicits darting behaviour in *P. maniculatus* but not *P. polionotus*. Together, we trace the evolution of species-specific escape thresholds to a central circuit node, downstream of peripheral sensory neurons, localizing an ecologically relevant behavioural difference to a specific region of the complex mammalian brain.

## INTRODUCTION

To survive in the wild, animals must respond to external sensory stimuli with actions appropriate for their local environment. Variation in behavioural responses may arise through learning, behavioural plasticity, or evolve through heritable changes of the underlying neural circuitry. For the latter, changes in sensory detection and/or processing have been shown to underlie behavioural evolution (e.g., host preference in mosquitos [McBride *et al*. 2014]; food preference in birds [Baldwin *et al*. 2014], cockroaches [Wada-Katsumata *et al*. 2013] and fruit flies [Auer *et al*. 2020]). When known, these sensory changes are most often due to genetic changes in peripheral sensory systems (e.g., odor or taste receptors, opsins [Tierney 1995; Cande *et al*. 2013], but see Seeholzer *et al*. 2018). By contrast, how evolution modifies central neural circuits to alter the innate behavioural responses of animals is less well understood (Roberts *et al*. 2022).

Visual stimuli have long been used to study defensive behaviours. Perhaps most famously, Tinbergen recorded the behaviour of birds exposed to cardboard models of aerial predators (Tinbergen 1948; Tinbergen 1951). This paradigm has since been modified to study naturalistic antipredator response to overhead visual stimuli under controlled conditions (Schiff *et al*. 1962; Ball and Tronick 1971; Holmqvist 1994; Hatsopoulos *et al*. 1995; Yamamoto *et al*. 2003; Yilmaz and Meister 2013; Temizer *et al*. 2015; De Franceschi *et al*. 2016). In this assay, laboratory mice (genus *Mus*) tend to freeze when exposed to a gliding overhead predator (“sweeping” stimulus), and most often flee or escape when exposed to an attacking predator (“looming” stimulus). Robust behavioural responses, such as these, have been used to uncover the underlying neural circuits, including a key role for the superior colliculus (SC) in translating visual stimuli into appropriate defensive reactions (Westby *et al*. 1990; Basso and May 2017; Wheatcroft *et al*. 2022), with projections from the retinorecipient superficial SC (sSC) to, for example, the deep layers of the SC (dSC) and on to the dorsal periaqueductal gray (dPAG; May 2006; Gale and Murphy 2014; Evans *et al*. 2018; Gale and Murphy 2018; Xie *et al*. 2021). Notably, dPAG neurons have been shown to command the initiation of escape actions (Lefler *et al*. 2020).

These defensive behaviours, and their underlying neural circuits, may diverge in species that have evolved in distinct environments, in which different defensive strategies may be more or less effective (Abrams 2000). Deer mice (genus *Peromyscus*) occupy diverse habitats across North America (Bedford and Hoekstra 2015), including species living in the underbrush of densely vegetated habitats (*P. maniculatus*) or those specialized for life in exposed, open fields with little to no vegetation (*P. polionotus*). Using these two ecologically divergent sister species, we show that they differ in behavioural response to the same visual threat and then identify a locus in the neural circuit where evolution has likely acted.

### Ecologically distinct *Peromyscus* species show contrasting defensive strategies

To test if defensive behaviours differ among mice from distinct habitats, we selected two closely related species of *Peromyscus*: the open field specialist *P. polionotus subgriseus*, and densely vegetated prairie inhabitant, *P. maniculatus bairdii.* A third species, *P. leucopus,* which is largely sympatric with *P. maniculatus*, was included as an outgroup to determine along which lineage any observed differences evolved (**Fig. 1A**). To quantify defensive behaviours, we placed wild-derived, laboratory-born adult mice in a large, open arena that included a refuge (i.e., a hut; **Fig. S1A)** and measured their response to an overhead “sweep-looming” stimulus, which resembles an aerial predator searching for (sweep), and then rapidly descending upon (loom), its prey. During the stimulus’s sweeping phase, mice from all three species generally decelerated and largely remained immobile (**Fig. 1B-C**; **Fig. S1B-C**; **Fig. S2A-B; Supplemental Movies 1-2**). Similarly, we did not observe any species-specific differences in behaviour when mice were exposed to either no stimulus (**Fig. S2C**) or an innocuous dimming stimulus (**Fig. S2D**). Conversely, the response to a looming stimulus revealed striking differences between species (**Fig. 1B-C**; **Fig. S1B-C; Supplemental Movies 1-2**). Both *P. maniculatus* and *P. leucopus* mice accelerated and ran rapidly across the arena (i.e., “darting”), often towards the refuge. By contrast, the open-field specialist, *P. polionotus,* tended to remain immobile (i.e., “pausing”). Phylogenetic comparison of these behaviours suggests that pausing in response to a looming stimulus is derived, and therefore the change in defensive response likely evolved along the *P. polionotus* lineage. Because the largest behavioural difference observed was in response to looming, we focused on this threat stimulus for subsequent experiments.

**Figure 1.**
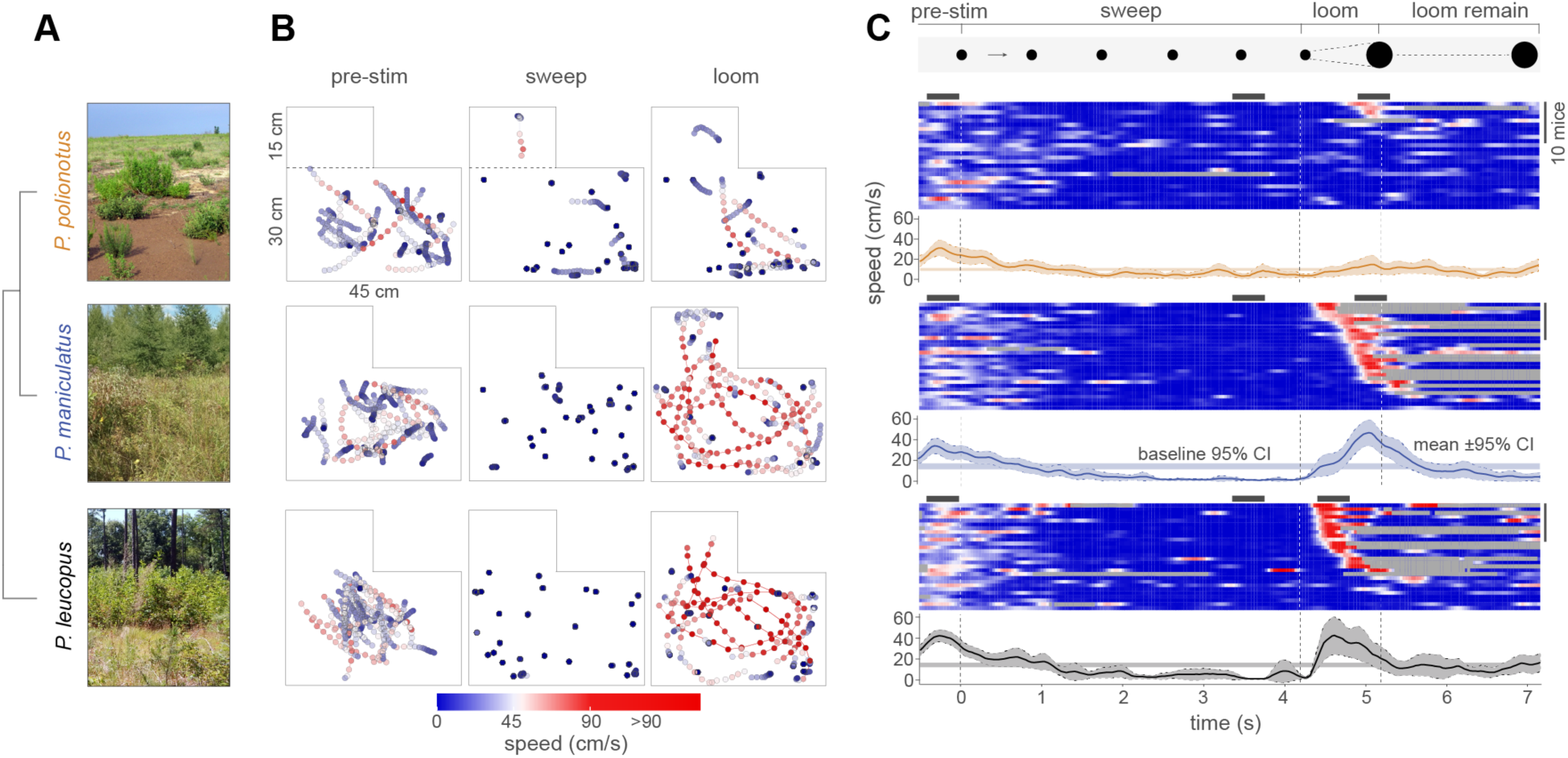
Evolution of defensive behaviour in ecologically distinct *Peromyscus* species. (**A**) Phylogenetic relationship of three focal *Peromyscus* species with representative photos of their natural habitat. (**B**) Movement trajectories of individual mice of *P. polionotus* (n=26), *P. maniculatus* (n=29) and *P. leucopus* (n=28) during 0.4 s before stimulus onset (left), sweeping (middle), and looming (right). Speed is indicated by a color gradient. (**C**) Raster plot of mouse speed during the sweep-looming stimulus (100% contrast). Rows represent individual mice. Trials are sorted by escape onset during the looming stimulus, with earliest on top. Speed color gradient is the same as in (B), with grey indicating the mouse is in the hut. Three grey bars above each raster plot indicate the time period of the trajectories shown in (B), and for looming are centered on the peak mean speed of each species. Line plots represent mean speed ± 95% confidence limit; horizontal shaded lines represent the 95% confidence interval of mean speed averaged across the 60 s before stimulus onset. Sample sizes are the same as (B).

### Escape threshold differences underlie the species-specific behaviours

With increasing threat intensity, prey animals often switch from immobility to rapid escape (De Franceschi *et al*. 2016; Branco and Redgrave 2020). To determine whether *P. maniculatus* and *P. polionotus* show similar changes in behaviour, we exposed a new cohort of mice from each species to five repetitions of a looming-only stimulus that varied in contrast (i.e., threat intensity; **Fig. 2A**; **Fig. S2–S3**). At low contrast (32%), most individuals of both species paused (14/20 *P. maniculatus*, 14/18 *P. polionotus*), while only a few mice darted (5/20 and 4/18, respectively). As the contrast level increased, the proportion of darting animals increased in both species, but the rate of change differed significantly between the species (**Fig 2A**). For example, at intermediate contrast (72%), most *P. maniculatus* (24/25) but few *P. polionotus* (4/23) darted, whereas, at high contrast (100%), the proportion of mice that darted was not significantly different between species (24/27 *P. maniculatus*, 19/27 *P. polionotus*). However, even at high contrast, the onset of darting was significantly delayed in *P. polionotus* (**Fig. 2B; Supplemental Movie 3**). Notably, contrast sensitivity curves obtained from single neuron recordings in the sSC did not differ between the two species, suggesting that a difference in stimulus detection does not explain the species-specific behavioural responses (**Fig. 2C**, **Fig. S4**). Together, we find that both species are more likely to pause at low and dart at high threat levels, but that the threat level (“threshold”) at which each species switches from pausing to darting differs: *P. maniculatus* transition to escape behaviour at an approximately two-fold lower threat intensity than *P. polionotus*.

**Figure 2.**
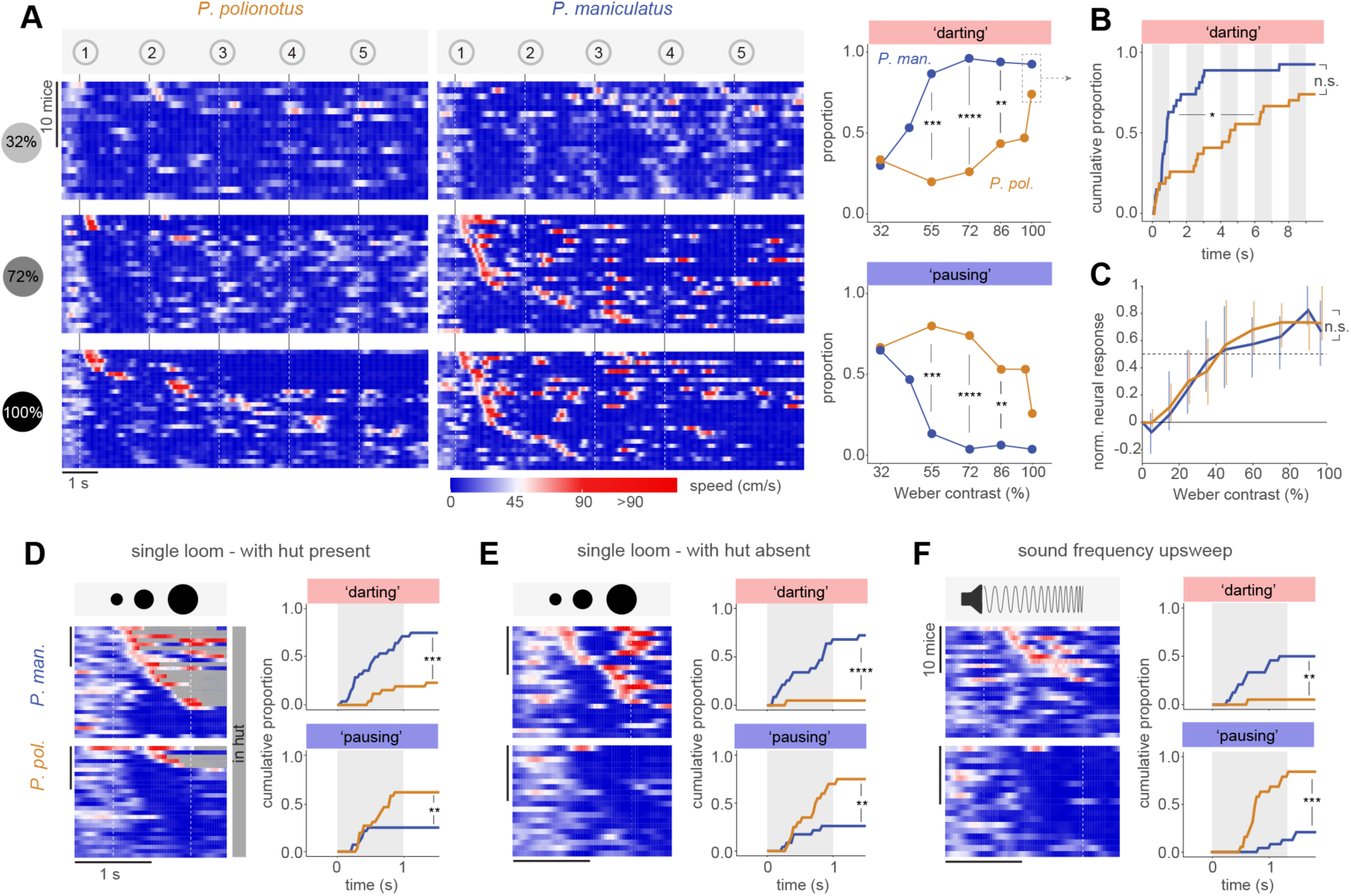
Escape threshold differences underlie species-specific behaviour. (**A**) Behavioural response to visual threat of varying intensity (looming contrast: 32%, 72%, 100%). Rows represent individual mice of *P. polionotus* (left) and *P. maniculatus* (right). Trials are sorted by latency to darting threshold. Proportion of individual mice of *P. polionotus* (gold) and *P. maniculatus* (blue) showing darting (top) and pausing (bottom) across these and additional contrast levels (far right). (**B**) Cumulative proportion of individual mice showing darting during 100% contrast looming stimulus. (**C**) Normalized peak firing rates of sSC neurons in head-fixed mice exposed to looming stimulus at different contrasts (n=3 animals for each species). Medians and 25-75% quantile ranges are indicated for each species. Firing rates are normalized to 0 for background firing and 1 for maximal response. (**D-F**) Raster plots and cumulative proportion of individual mice showing darting and pausing during a single looming stimulus (100% contrast) in the presence of hut (D), in the absence of hut (E), and during a sound frequency upsweep (F). Chi-Squared test (proportion, cumulative proportion), Kolmogorov-Smirnov test (darting onset distribution), Wilcoxon rank-sum test (firing rate). n.s. not significant; * *P* < 0.05; ** *P* < 0.01; *** *P* < 0.001; **** *P* < 0.0001.

We next asked if spatial context and/or stimulus modality affect the observed differences in defensive behaviour. First, we compared the response to the looming stimulus of two new cohorts of mice that either had access to a refuge or not. In the absence of a hut, the species-specific responses were recapitulated (darting with hut: 21/28 *P. maniculatus*, 6/26 *P. polionotus*; darting without hut: 17/23 *P. maniculatus*, 1/20 *P. polionotus*) (**Fig. 2D-E; Supplemental Movies 4-5),** suggesting that differences in the perception of safety afforded by the refuge is not driving the species-specific response. To determine if the observed behavioural differences are specific to visual stimuli, we exposed a new cohort of mice to an aversive ultrasound frequency upsweep (Mongeau *et al*. 2003; Vale *et al*. 2017). Again, we observed remarkably similar species-specific behaviour: many *P. maniculatus* accelerated and some darted (12/24 darted, 5/24 paused), while *P. polionotus* primarily paused (1/19 darted; 15/19 paused) (**Fig. 2F; Supplemental Movies 6-7**). Collectively, these data suggest that the species-specific behaviour is consistent with context- and modality-independent differences in escape threshold.

### Differential activation of dPAG during escape behaviour

We next sought to identify the neural circuit component(s) where evolution acted to generate the observed differences in defensive behaviour. Our electrophysiological recordings in head-fixed mice suggest that visual threat information is faithfully relayed to the retinorecipient sSC in both species (**Fig. 2C**). Moreover, we found that an aversive auditory stimulus can recapitulate the visually triggered behaviour, suggesting the neural mechanism is likely located downstream of visual and auditory inputs (**Fig. 2F**). Because the medial dSC and dPAG play a central role in mediating escape behaviours in response to both visual and auditory stimuli in rodents (Sahibzada *et al*. 1986; Coimbra *et al*. 1989; Vargas *et al*. 2000; Bittencourt *et al*. 2005; Shang *et al*. 2015; Wei *et al*. 2015; Deng *et al*. 2016; Evans *et al*. 2018), we hypothesized that differences in the recruitment of the dSC-dPAG pathway could explain the species-specific responses at the behavioural level.

To test this hypothesis, we characterized neural activation in the dSC and dPAG during a darting response, using the immediate-early gene c-Fos as a proxy (Bullitt 1990). We first confirmed that prolonged exposure to high contrast looming stimuli triggered repeated darting in both species (as in Fig. 2). After dark adaptation, we exposed individuals to 25 sets of 5 looming stimuli and recorded their behavioural responses (**Fig. 3A**). As expected, *P. maniculatus* darted more often and faster than *P. polionotus*, but darting mice (with more and faster darts than control animals) could be identified in both species (**Fig. 3B**). In a subset of these mice (**Fig. S5A**), representing the species-typical responses, we counted the number of c-Fos+ cells (**Fig. 3C**). First, in looming-exposed mice of both species, we found higher numbers of c-Fos+ cells, compared to control mice, in the medial dSC, but not the lateral dSC, which view the upper and lower visual field, respectively (DrÄger and Hubel 1975), consistent with the overhead position of the looming stimulus (**Fig. 3D**; **Fig. S5B-E**). In the medial dSC, the number of c-Fos+ cells correlated with mean speed during darting but not species identity (**Fig. S5F**). Thus, the dSC was active in looming-exposed mice, but levels of neural activation did not differ between species.

**Figure 3.**
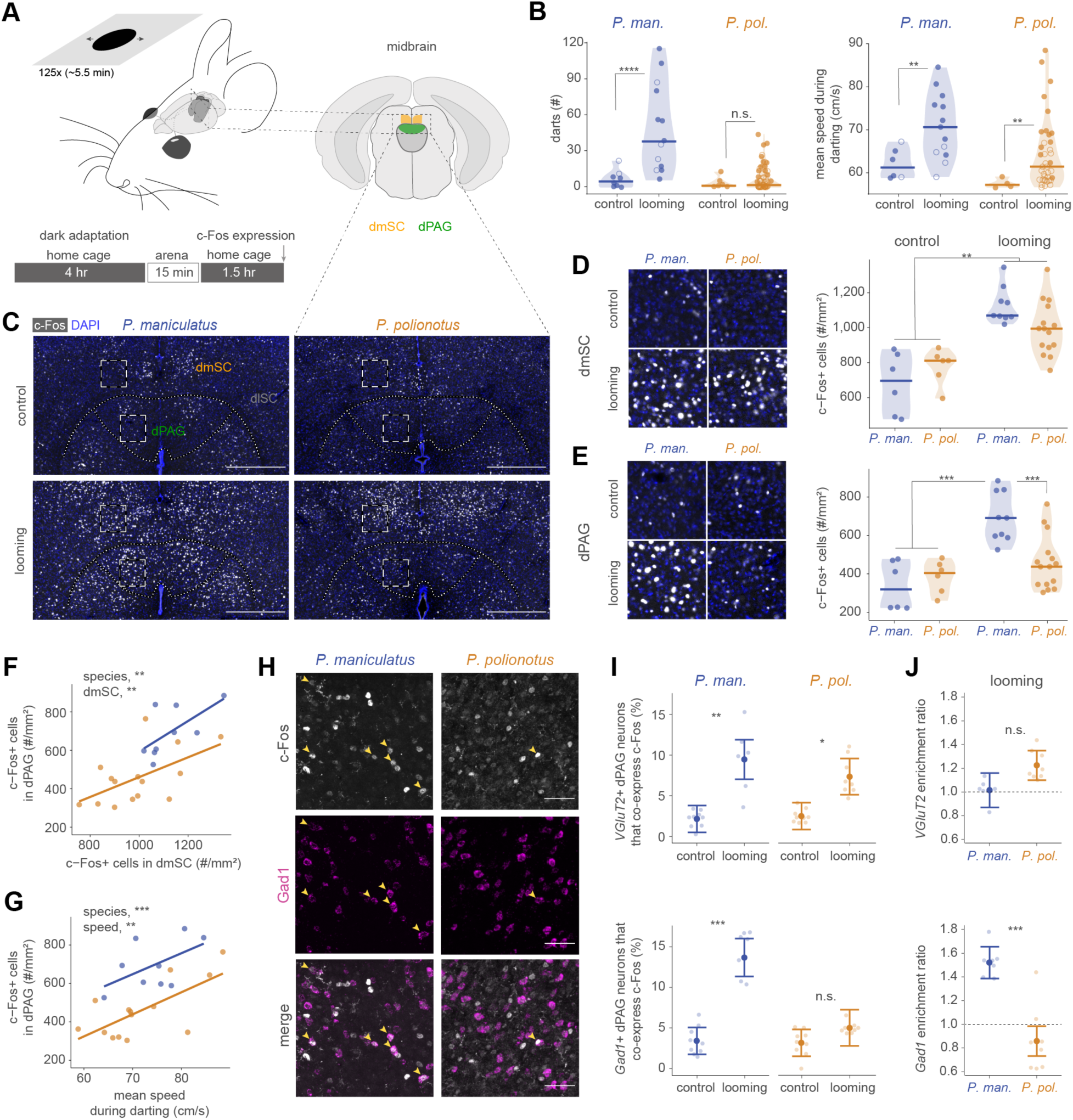
Differential activation of dPAG neurons during escape behaviour. (**A**) Behavioural setup and schematic of assay to measure neural activation during escape. Two focal midbrain regions are shown: dmSC (orange) and dPAG (green). (**B**) Number of darts (left) and mean speed during darting (right) of *P. maniculatus* (blue) and *P. polionotus* (gold). Data from mice that were included in the c-Fos experiment are indicated with filled circles (looming, *P. maniculatus*, n=9; *P. polionotus*, n=15; control, n=6 for both species). (**C**) Representative images of c-Fos expression in dmSC and dPAG. Scale bar, 500 μm. Dashed white boxes indicate regions that are enlarged in (D-E). (**D-E**) Images (left) and quantification (right) of c-Fos+ cells in the dmSC (D) and dPAG (E) of control and looming-exposed mice. Sample sizes are reported in (B). (**F**) Quantification of c-Fos+ cells in dPAG as a function of number of c-Fos+ cells in dmSC of looming-exposed mice. (**G**) Number of c-Fos+ cells in dPAG as a function of mean speed during darting in looming-exposed mice. (**H**) Representative images of c-Fos (top), Gad1 (middle) and merged (bottom) staining in the dPAG. Yellow arrows indicate positively (or double) stained cells. Scale bar, 50 μm. (**I**) Proportion of c-Fos+ excitatory (VGluT2; top) and inhibitory (Gad1; bottom) dPAG neurons in control and strongly escaping mice. Model fit and 95% confidence interval are shown. Points represent tissue sections (n=6-9 sections from 3 mice per species). (**J**) Enrichment index [proportion of excitatory/inhibitory neurons that co-express c-Fos divided by the overall proportion of c-Fos+ neurons] for excitatory (top) and inhibitory (bottom) neurons in dPAG of strongly escaping mice. Statistical significance evaluated with mixed effects models. n.s. not significant; * *P* < 0.05; ** *P* < 0.01; *** *P* < 0.001; **** *P* < 0.0001.

In contrast, the dPAG showed species-specific differences in neural activation. Overall, the number of c-Fos+ cells was high in looming-exposed *P. maniculatus*, but low in *P. polionotus,* similar to control animals (**Fig. 3E**; **Fig. S5G-H**). Variation in dPAG activation across looming-exposed mice correlated with dSC activation and mean speed during darting, but not with the number of darts (**Fig. 3F-G**; **Fig. S5I**; see also **Fig. S6**). However, dPAG activation in *P. maniculatus* was consistently ~1.5-fold higher compared to *P. polionotus* across dSC activation levels or darting speeds (**Fig. 3F-G**). Thus, exposure to visual threat and resulting escape movement does not lead to an increase in c-Fos+ levels in the dPAG of *P. polionotus* to the same extent as in *P. maniculatus*.

In *Mus*, excitatory neurons in the dPAG can initiate rapid escape (Deng *et al*. 2016; Evans *et al*. 2018). Using single-molecule fluorescent *in situ* hybridization, we examined the transmitter identity of c-Fos+ neurons in mice with the strongest escape responses and c-Fos+ levels (**Fig. 3H**). We found that both species possess comparable numbers of excitatory and inhibitory neurons in the dSC and dPAG, and both classes are activated in the dSC of the same animals during visually evoked escape (**Fig. S7**). However, while excitatory dPAG neurons were activated in both species, a greater number of c-Fos+ inhibitory neurons was detected in the dPAG of looming-exposed *P. maniculatus,* but not in *P. polionotus,* compared to control animals (**Fig. 3I**). In addition, inhibitory neurons in looming-exposed *P. maniculatus* were approximately 1.5-fold more frequently activated than dPAG neurons in general (**Fig. 3J**). Together, these immunohistochemistry results suggest that visually-evoked darting in *P. polionotus* does not recruit the same ensemble of dPAG neurons that is activated in *P. maniculatus*.

To measure neural activity directly and to disentangle the effects of visual threat exposure and behavioural response, we next used Neuropixels probes to record from the SC and dPAG in head-fixed mice that were running on a spherical treadmill and intermittently exposed to looming stimuli (**Fig. 4A and S8**). Visual responses to looming stimuli were found in the SC and dPAG of both species (**Fig. 4B**), and both the neural responses (**Fig. 4C and S9**) and behavioural reactions (**Fig. S2D**) to looming were stronger than responses to an innocuous dimming stimulus, consistent with results in *Mus* (Yilmaz and Meister 2013; Lee *et al*. 2020). To explore the relationship of dPAG activity and motor behaviour, we examined time periods when mice initiated running, in the absence of visual stimulation, after a period of slow walking or immobility (**Fig. 4D**, top row). Single neurons (**Fig. 4D** bottom) as well as the population of dPAG neurons (**Fig. 4E**) increased their activity during these running periods in *P. maniculatus* but not in *P. polionotus*. Consistent with results from immunohistochemistry, neural activity correlated with movement only in *P. maniculatus* (**Fig. 4F**; **Fig. S10**). Furthermore, a comparison of running speed and neural firing patterns revealed that neural activity in the dPAG of *P. maniculatus* precedes the onset of running (**Fig. 4G**; **Fig. S10**).

**Figure 4.**
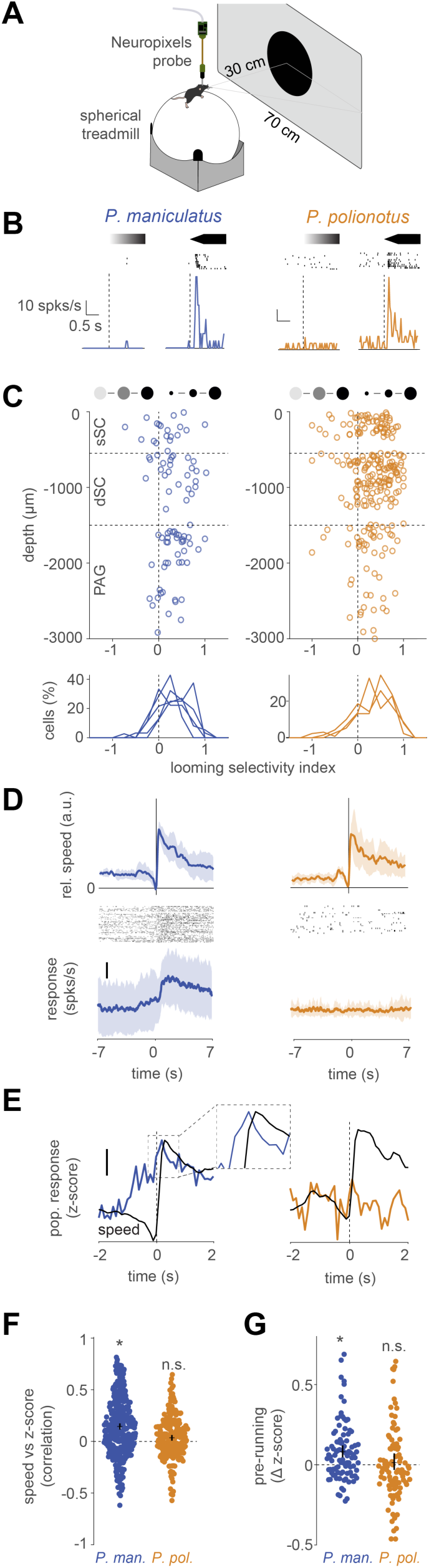
Species-specific encoding of locomotion in the dPAG. (**A**) Schematic of setup to record neural activity and running speed. Mice were head-fixed on a floating ball and presented with looming/dimming stimuli. (**B**) Representative responses to dimming (left) and looming (right) stimuli of one cell in the dPAG for *P. maniculatus* (blue) and *P. polionotus* (gold). (**C**) Looming selectivity of neurons in the SC and PAG for looming and dimming stimuli. Indices for sSC, dSC and dPAG are similar for both species (*P* > 0.1 for each, two-sample Kolmogorov-Smirnov test). Bottom: Distributions indicate looming selectivity indices for each tested mouse (*P. maniculatus,* n=4; *P. polionotus,* n=3). Dashed vertical line represents selectivity index = 0. (**D**) Speed and neural activity during onset of running. Mean ± STD of running speed (top row) around running onset (n=3 animals for each species). Raster plot (middle row) and mean ± STD (bottom row) of an example neuron in the dPAG. Scale bar: 10 spks/s. (**E**) Mean z-score of all dPAG neurons in all mice (*P. maniculatus*, blue; *P. polionotus*, gold), overlayed with the speed trace (black). Scale bar: 0.1. Inset shows neural activation precedes behaviour. (**F**) Correlation coefficient of speed with z-score of mean neural activity in the dPAG. Mean (horizontal line) and estimated 95% confidence intervals (vertical line) are shown relative to zero (dashed line) (see also Fig. S10C). (**G**) Difference in z-score in 600 ms before running onset. Mean (horizontal line) and estimated 95% confidence intervals (vertical line) are shown relative to zero (dashed line) (see also Fig. S10D). * *P* < 0.05 unpaired median difference Gardner-Altman estimation.

Taken together, results from immunohistochemistry and electrophysiology indicate that visual threat can trigger neural activity in the dSC and dPAG of both species. However, while dPAG neurons encode running events and darting in *P. maniculatus*, dPAG activity does not correlate with the initiation of running or escape behaviour in *P. polionotus*, suggesting the dPAG may contain the neural circuits on which evolution has acted.

### Optogenetic activation of excitatory dPAG neurons recapitulates species differences

To investigate the causal role of the dPAG in mediating behaviours, we optogenetically activated neurons in the dPAG of both *P. maniculatus* and *P. polionotus*. An AAV2 vector was injected bilaterally into the dPAG to express channelrhodopsin in excitatory neurons, under the control of the CamKII promoter (**Fig 5A**; **Fig. S11**, **Fig. S12**). Using a centrally implanted optic fiber (**Fig. 5B**), we stimulated the dPAG as mice moved freely in a circular arena. The trajectory and speed of mice was extracted, and each trial was classified as forward movement, slowing or “other” based on the behaviour of the animal during the stimulation (**Fig. 5C-E**, **Fig. S12**, **Suppl. Movies 8-12**; see Methods). Similar to our observations during visual stimulation, we found that while *P. maniculatus* exhibited more forward movement during the optogenetic stimulation compared to controls, *P. polionotus* did not (**Fig. 5F**). Instead, *P. polionotus* tended to slow or pause during stimulation (**Fig. 5G**). These species-specific differences in evoked behaviours were also evident when comparing the distribution of speed change (**Fig. 5H**, **Fig. S13**). Consistent with c-Fos+ levels (**Fig 3**), increasing laser power during optogenetic stimulation triggered faster forward movement events in *P. maniculatus*, while the speed of both forward accelerations and slowing events decreased in *P. polionotus* (**Fig. 5I-J** and **S13**). These findings, together with the c-Fos analysis and *in vivo* electrophysiological recordings, demonstrate that while the dPAG plays an important role in mediating forward acceleration, including running behaviours, in P. *maniculatus*, it does not play the same role in *P. polionotus*.

**Figure 5.**
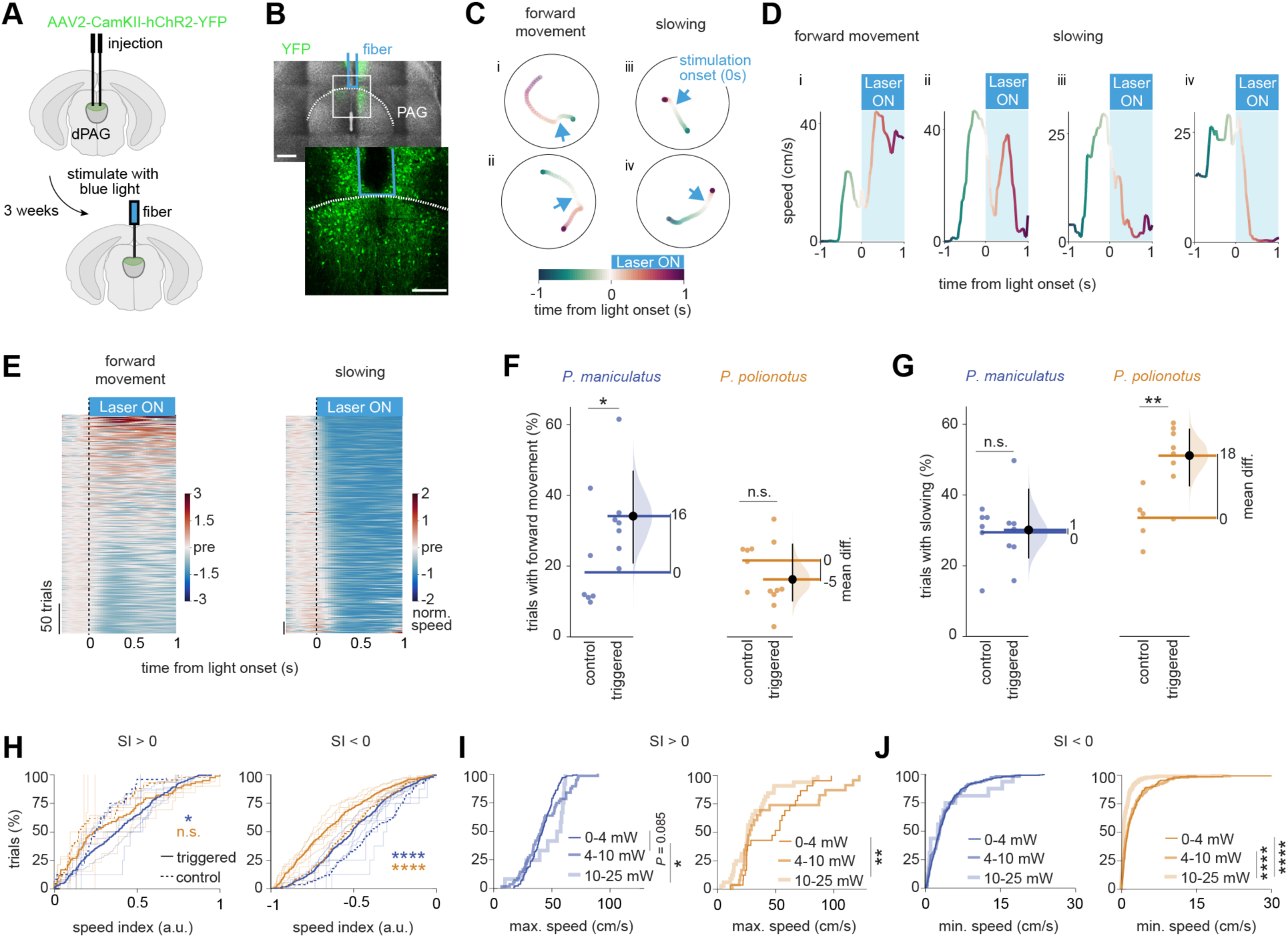
Optogenetic activation of excitatory dPAG neurons recapitulates species differences. (**A**) Experimental paradigm for optogenetic activation of the dPAG. (**B**) YFP-ChR2+ neurons (green) and optic fiber tract (blue) in the dPAG. Scale bar: 400 μm (top image), 200 μm (bottom image). (**C**) Example trajectories of individual mice showing forward movement (i and ii; see Suppl. Movie 8 and 9) and slowing (iii and iv; see Suppl. Movie 10 and 11). Color indicates time before (greens) and during (reds) laser stimulation. (**D**) Speed traces of mice corresponding to example trajectories in C. (**E**) Raster plot of normalized speed of all trials classified as ‘forward movement’ (left) or ‘slowing’ (right). Speed was normalized to the 0.37 s before laser stimulation (‘pre’) and is indicated by a color gradient. (**F-G**) Percentage of trials with forward movement (F) and slowing (G) behaviour for *P. maniculatus* (left; triggered, n = 7; control, n = 6) and *P. polionotus* (right; triggered, n = 8; control, n = 5). Means (horizontal lines) for control and triggered mice and estimation statistics (unpaired mean difference Gardner-Altman estimation) are shown; distribution represents 5000 bootstrapped samples. Mean difference (black dot) and confidence intervals (vertical black line) are provided. (**H**) Cumulative distribution of speed index (SI; see Suppl. Fig. 13) for SI>0 (left) and SI<0 (right) for *P. maniculatus* (blue; triggered: n = 706 trials [n = 7 mice], control: n = 138 [n = 6]) and *P. polionotus* (gold; triggered: n = 568 trials [n = 8 mice], control: n = 215 [n = 5]). Individual mice are shown as thin lines; cumulative distributions of all trials as thick lines (triggered) or thick dashed lines (control). (**I**) Cumulative distribution of maximum speeds for trials with SI>0 at three different ranges of laser power (0-4, 4-10, 10-25 mW) for *P. maniculatus* (blue) and *P. polionotus* (gold), separately. (**J**) Cumulative distribution of minimum speeds for trials with SI<0. * *P* < 0.05, ** *P* < 0.01, **** *P* < 0.0001, two-sample Kolmogorov-Smirnov test.

## DISCUSSION

Here we show that ecologically distinct, yet closely related, species of deer mice evolved species-specific differences in defensive behaviour. Specifically, when exposed to a looming stimulus, *P. polionotus* requires a higher threat intensity to reliably trigger a darting response relative to both *P. maniculatus* and *P. leucopus*.

It is possible that the observed species-specific behaviour evolved via natural selection. In dense fields or forests, escape (e.g., darting to a refuge) may increase survival probability, whereas in open environments, with fewer refuges, movement may be conspicuous and pausing/freezing could minimize predator detection. However, when exposed to an intense threat (i.e., imminent attack), escape may be the only survival option. This explanation is consistent with the observation that *Mus* are more likely to freeze in the absence of a refuge (Vale *et al*. 2017), and more broadly with the distinct strategies observed in other species to avoid predator detection in different environments (Wywialowski 1987; Lima and Dill 1990; Wheatley *et al*. 2020).

A central question remains: where are these ecologically relevant behavioural differences encoded in the brain? Two observations suggest that the neural mechanism is located centrally. First, visual looming stimuli trigger similar patterns of neural activity across the entire depth of the SC. Second, both visual and auditory stimuli evoked similar species-specific behavioural responses. Together, these data indicate that the key neural mechanism lies downstream of both the retina and the confluence of sensory inputs in the dSC. This is different from the many studies (Seehausen *et al*. 2008; Montgomery *et al*. 2021; Lloyd *et al*. 2022) that link behavioural evolution to peripheral sensory systems. Indeed, we find differences – in both neural activity and optogenetic manipulation – that point to the dPAG, a subregion of the PAG specifically implicated in driving escape behaviour (Branco and Redgrave 2020), as the likely locus of evolution.

There are several hypotheses for how the dPAG may differ between *Peromyscus* species. First, the dPAG may receive different input (e.g., in the number or strength of projections from the SC). However, we found that the dPAG is similarly activated in both species when animals were head-fixed and exposed to a visual looming stimulus, suggesting the dPAG receives the same input from the SC. Second, the two species may differ in the number of excitatory cells in the dPAG; in *Mus*, activation of excitatory cells leads to an escape response (Evans *et al*. 2018). However, we do not find a measurable difference in the number of excitatory (or inhibitory) neurons in the dSC or dPAG. Third, the properties of excitatory neurons or their circuit connections may be different in the two species. For example, the intrinsic excitability of neurons could differ (Burnett *et al*. 2022), or be held at different levels through long-range inhibitory projections from other brain regions (Tovote *et al*. 2016; Fadok *et al*. 2017; Fratzl *et al*. 2021; Salay and Huberman 2021; Li *et al*. 2022). Alternatively, the role of the dPAG in the circuit could be reversed, for example, through changes in local wiring or in long-range projections, such that exposure to the same threat triggers distinct neural computations and defensive responses. Future work is aimed at determining the molecular mechanisms that explain why incoming threat information is not sufficient to engage the ensemble of excitatory and inhibitory dPAG neurons in *P. polionotus,* unlike *P. maniculatus*, to initiate escape behaviour.

Here we show that ecologically distinct deer mice evolved species-specific defensive behaviours and trace this difference to a central brain region, the dPAG. Together our data suggest that evolution can tinker with a behaviour dial by shifting the threshold between two conserved behaviours – pausing and darting – to fine-tune defensive response in different environments, providing a rare example of a central brain region linked to natural variation in a sensory-driven behaviour.

## ACKNOWLEDGEMENTS

We thank Melis Yilmaz, Markus Meister, Max Joesch, and Tiago Branco for advice on the behavioural experiments; Catherine Dulac, Vassilis Bitsikas, Erin Diel, and Jenny Chen for advice on the immunohistochemistry and RNAscope experiments; Joel Greenwood and Ed Soucy for technical and engineering help; Arnau Sans Dublanc and Anna Chrzanowska for help and advice on optogenetic experiments; Bram Nuttin for help with DeepLabCut; Steven Worthington for statistical advice; and Renate Hellmiss for figure design. Bernardo Sabatini, Vanessa Stempel, Kelsey Tyssowski, and Nacho Sanguinetti gave helpful feedback on the manuscript. Yu Man Lee and Aimee Tomcho kindly provided the habitat pictures of *P. maniculatus* and *P. leucopus*, respectively, in Fig. 1. Brain silhouettes in Figs. 3 and 5 were adapted from the Allen Mouse Brain Reference Atlas. FB was supported by an HHMI International Student Research Fellowship, a Grant-in-Aid of the American Society of Mammalogy, a Herchel Smith Graduate Fellowship, a Robert A. Chapman Memorial Scholarship, and a Joan Brockman Williamson Fellowship. This project has received funding from the European Union’s Horizon 2020 research and innovation programme under the Marie Skłodowska-Curie grant agreement No 665501 and by the FWO (12S7917N and 12S7920N) to KR. VT was supported by a Harvard PRISE fellowship and a Harvard Museum of Comparative Zoology grant for undergraduate research. KF is supported by the FWO (G094616N and G091719N) and the NIH (1R01EY032101). HEH is an Investigator of the Howard Hughes Medical Institute.

## MATERIALS & CORRESPONDENCE

Correspondence to Hopi E. Hoekstra, Felix Baier, or Karl Farrow.

## DATA AVAILABILITY

All relevant source data will be included upon publication.

## CODE AVAILABILITY

The code used for the analyses will be available from GitHub.

## MATERIAL & METHODS

### Mouse strains & husbandry

Colony founders of *P. maniculatus bairdii* (strain BW), *P. polionotus subgriseus* (strain PO), and *P. leucopus* (strain LL) were originally obtained from the *Peromyscus* Genetic Stock Center at the University of South Carolina and then established and maintained at Harvard University. Behaviour, c-Fos and RNAscope experiments were performed at Harvard and, later, optogenetics and *in vivo* recording experiments at Neuro-Electronics Research Flanders (NERF).

Housing at Harvard University: We housed all animals on Bed-o’Cobs 1/4” bedding (The Andersons, Maumee, Ohio) in ventilated standard rodent cages (Allentown Inc., Allentown, NJ) on a 16 hr light: 8 hr dark cycle at 23°C. We provided animals with a red translucent polycarbonate hut, Enviro-Dri nesting material, and a cotton nestlet. All animals were given *ad libitum* access to irradiated Prolab Isopro RMH 3000 5P74 (LabDiet) and water.

Housing at NERF: Animals were housed on Lignocel 3-4 bedding (J. Rettenmaier & Söhne GmbH, Germany) in ventilated standard rodent cages on a 12 hr light:12 hr dark cycle at 23°C. Animals were provided with cotton nesting material. All animals were given *ad libitum* access to chow diet (ssniff, Soest, Germany) and water.

After weaning litters at 23 days of age, we kept same sex animals in groups of up to five individuals by strain, unless otherwise indicated.

### Behaviour experiments

#### Experimental setup

To assay behavioural response to a visual stimulus, we constructed a rectangular behavioural arena from plexiglass that measured 45 cm (W) × 30 cm (D) × 30 cm (H), adapted from (Yilmaz and Meister 2013). We attached a triangular prism-shaped hut (24 cm [W] × 18 cm [D] × 12 cm [H]) to one corner of arena floor. To reduce reflection, we covered the arena walls and floor with Matte Finish (Krylon). To illuminate the arena, we lined the outside base of the walls with infrared light (IR) LED strips. To record behaviour from below the arena, we made the ground floor of IR-transmissive black plexiglass and used an IR-sensitive camera (Flea3 FL3-FW, monochrome, Point Grey Research) to record at 30 fps. We programmed visual stimuli with Psychtoolbox-3 for Matlab (Brainard 1997; Kleiner *et al*. 2007) and displayed them on an LCD monitor from above the arena. Finally, we triggered an LED (invisible to the mouse) simultaneous to the visual stimulus through an Arduino Uno connected to the computer, which we used to synchronize individual frames with the stimulus. We generated sound stimuli with a power amplifier (TB10A, Fosi Audio) connected to a tweeter (Pro-TW120, DS18).

#### Experimental procedure

Before each behavioural experiment, animals were left undisturbed in their cage for 24 hrs. We conducted all experiments within the first 4 hrs of the dark period (Zeitgeber time) and in red light. We habituated animals to the experiment room for 30 mins, and then transferred a single individual to the behavioural arena, where it habituated for 10 mins. We triggered the stimulus manually when the mouse moved away from the walls towards the center of the arena and recorded the behaviour of the mouse for 2-3 mins before and after the stimulus was triggered. Once testing was complete, we moved the individual to an empty cage and wiped out the arena with 70% ethanol. We then assayed the remaining individuals in the cage following the same protocol.

For the contrast experiment, we randomly assigned a new cohort of mice and exposed each mouse once to a contrast level. Approximately 1 week (range of 5-11 days) after the first exposure, we again randomly assigned the same individuals to a different contrast level and exposed them once. We employed this approach to both minimize habituation from repeated testing and to reduce the number of animals needed for the experiment.

For all other experiments (sweep-looming, looming with hut, looming without hut, dimming, auditory), we used new cohorts of naïve animals and exposed individuals to the stimulus only once.

To determine which brain region(s) show activity correlated with behavioural response, we collected the brains of animals following their exposure to the overhead stimulus. To this end, we single-housed animals in a new cage the day before the experiment. On the test day, we dark habituated the animals by moving their cage in the test room for 4 hrs. We then gently transferred animals to the arena, with the hut closed off. After 10 mins of habituation, we triggered the stimulus. We recorded the behaviour of animals during the complete trial. We then transferred animals back into the cage and transcardially perfused them after 90 mins in the dark (see below).

#### Stimuli

To quantify response to a visual stimulus, we first conducted an assay with a combined sweep-looming stimulus (De Franceschi *et al*. 2016). The stimulus was a black disc on gray background with a diameter of ~4° visual angle (approximately 2.2 cm) that first appeared in one corner of the computer screen and slowly moved diagonally at a speed of 10°/s. Once the disc reached the center of the screen, it rapidly expanded to a diameter of 40° visual angle (approximately 22 cm) at a linear speed of 36°/s. The disc then remained at full diameter for 2 s before disappearing. We chose these parameters because preliminary experiments revealed that they maximized the difference in behavioural response between the two focal species.

To measure the behavioural response of animals to different levels of threat, we altered the contrast of the looming disc by changing its intensity against the standard gray background. Intensity is indicated as a positive percentage, converted from the negative Weber fraction (Evans *et al*. 2018). We used different contrast levels of the looming disc: 32%, 55%, 72%, 86%, 100%, with one additional contrast level for each species within its dynamic range (*P. maniculatus*: 45%; *P. polionotus*: 97%). Contrast values were validated with a digital illuminance meter (LX1330B, Dr. Meter). The stimulus is comprised of 5 repeats of the standard looming stimulus, with a remain time at full diameter of 0.5 s and an inter-stimulus period of 0.5 s (Evans *et al*. 2018).

To test the behavioural response to an aversive auditory stimulus, we exposed animals in the looming arena to an ultrasound frequency upsweep (17-20 kHZ over 1.3 seconds, repeated five times; 80 dB at arena floor), while the visual screen displayed a gray background (Mongeau *et al*. 2003; Vale *et al*. 2017).

To test the effect of a refuge on behavioural response, we exposed animals to a single looming stimulus (black disc on gray background) in the presence of the hut. As before, the stimulus remained at full diameter for 2 s before disappearing. To test the effect of the absence of a refuge, we closed off the hut and exposed animals to the same single looming stimulus.

To test the behavioural response to a non-moving, innocuous visual stimulus, we used a disc of fixed size (diameter of 40°) that appeared in the center of the screen, initially matching the gray background but then changing to black over 1 s and remaining black for 2 s before disappearing.

To quantify c-Fos levels after defensive behaviour, we exposed animals to 125 repeats of the standard looming stimulus, structured into 25 sets of 5 repeats, with a remain time at full diameter of 0.5 s, an inter-stimulus period of 0.5 s within sets, and an inter-set period of 3 s. Control animals were exposed to only the standard gray background.

#### Analysis

To characterize the behavioural response of an animal to the stimulus, we used a custom Matlab code to retrieve centroid coordinates of the animal and the status of the stimulus from the video recordings. We calculated the speed of each animal from these coordinates and smoothed the data using a mean filter with a width of 5 frames.

We defined and automatically annotated “darting” behaviour as a speed ≥ 56 cm/s, and “pausing” behaviour as a speed of ≤ 3.28 cm/s for at least 0.4 s while the animal was outside the hut (see **Fig. S2**). We arrived at these definitions by comparing behaviour during exposure to a single looming stimulus to baseline behaviour. Specifically, we analysed a video segment for each animal with a duration of 1 s that preceded stimulus exposure by 1-2 mins. We selected video segments that matched our criterion for triggering a stimulus (see above; i.e., when the animal moved away from the walls towards the center of the arena). We recorded escape speed as the maximum speed during the darting event.

For the sweep-looming experiment, trials in which the animal was in the hut at the onset of the looming stimulus were removed (*P. maniculatus*, N=1; *P. polionotus*, N=3; *P. leucopus*, N=8); we compared only animals that were exposed to the full stimulus.

For the contrast looming experiment, we removed trials for which two independent observers did not unanimously confirm a discernible response (i.e., interruption or commencement of body movement) during the first looming repeat from the dataset (*P. maniculatus*, N=18; *P. polionotus*, N=50) to compare only animals that detected the stimulus. We detected a significant effect of both species and contrast level on the probability of a discernible response (logistic regression; species: *z* = −4.65, *P* = 3.36*e^−6^; contrast level, *z* = 5.56, *P* = 2.75*e^−8^; see **Fig. S3A** for % discernible response by contrast for each species).

To test for the effect of the presence/absence of a hut, we removed animals that did not show evidence of detecting the stimulus (hut present: *P. maniculatus*, N=1; *P. polionotus*, N=1; hut absent: *P. maniculatus*, N=0; *P. polionotus*, N=2).

### c-Fos experiments

#### Immunohistochemistry and imaging

To measure neuronal activity of animals exposed to a looming stimulus, we used the immediate early gene, c-Fos, as a marker of neuronal activity. Following the behaviour experiment described above, we transcardially perfused mice with ice-cold 1x phosphate-buffered saline and then with 4% paraformaldehyde. Brains were dissected out, postfixed for 24 hrs at 4°C, cryopreserved in 30% sucrose, and stored at −70°C until subsequent use. To stain for c-Fos protein, we sectioned brains at 40 μm, blocked tissue for 1 hr, and incubated sections for 2 days with rabbit anti-c-Fos antibody (1:4000, Synaptic Systems, 226003). We used donkey anti-rabbit Alexa 647 antibody (1:1000, Invitrogen, A31573) for secondary detection and mounted tissues with DAPI Fluoromount-G (SouthernBiotech, 0100-20). Slides were imaged on an AxioScan.Z1 slide scanner (Zeiss).

#### Analysis

Following imaging, we exported images to .tif format and arranged sections into anterior-posterior order with the ImageJ plugin TrakEM2 (Cardona *et al*. 2012). We manually outlined ROIs with custom Fiji (Schindelin *et al*. 2012) macros based on landmark structures identified using autofluorescence patterns and DAPI staining. To segment images, we used the ImageJ plugin StarDist (Schmidt *et al*. 2018) with default parameters (model – versatile, normalize image – yes, percentile low – 1, percentile high – 99.8, probability – 0.5, overlap threshold – 0.4), which automatically detects cells using neural network models with star-convex shape priors. For each identified cell in the dataset, we retrieved the area, X/Y coordinates, and mean intensity. We filtered out large artefacts that were incorrectly identified as cells by removing objects with an area of > 180 μm^2^. For each ROI and section separately, we then counted cells with mean intensities larger than the mode of the density function of mean intensities as c-Fos-positive. We chose this approach to remove cells with low c-Fos expression and to make cell counts robust against batch effects (e.g., different baseline c-Fos expression levels across experiments).

### Single-molecule fluorescent *in situ* hybridization (smFISH)

#### Experimental procedure

To determine if c-Fos+ cells were excitatory or inhibitory neurons, we selected, from our previous c-Fos experiment, three individuals of each species that showed the most extreme darting behaviour in response to the 5 min looming stimulus as well as three control animals for combined smFISH-IHC processing. We obtained six sections (thickness 14 μm) from each animal, and then used half to detect *NeuN*, *Gad1*, and c-Fos, and the other half to detect *NeuN*, *VGluT2*, and c-Fos. Seven of the 72 sections did not have reliable staining and were excluded from the dataset. To determine if AAV2+ cells were primarily excitatory or inhibitory, we injected three animals of either species with the viral vector (see ‘Virus injection and fiber implantation’ under ‘Optogenetic activation experiments’) and then obtained six sections (thickness 14 μm) from each animal and used half to detect *Gad1* and YFP, and the other half to detect *VGluT2* and YFP.

#### smFISH protocol

We used the RNAscope Multiplex Fluorescent Reagent Kit v2 with the RNA-Protein Co-Detection Ancillary Kit for co-detection of mRNA and protein. For smFISH, we used custom-made RNAscope probes for *Gad1*, *VGluT2* (*Slc17a6*), and *NeuN* (*Rbfox3*). Probes were based on the coding sequence of each gene, and single-nucleotide polymorphisms were included by alternating between species (*P. maniculatus*, *P. polionotus*). For IHC, we used the rabbit anti-c-Fos (1:100, Synaptic Systems, 226003) and rabbit anti-GFP (1:100, Thermo Fisher, A-11122) antibodies to detect c-Fos protein and the YFP tag in the viral vector, respectively, and HRP-labeled goat anti-rabbit antibody (1:500, PerkinElmer, NEF812001EA) for secondary detection. We visualized RNA probes and antibodies with Opal 520, Opal 570, and Opal 690 dyes (1:1000, Akoya Biosciences, FP1487001KT, FP1488001KT, FP1497001KT), and counterstained with DAPI. Regions of interest (mSC, dPAG) were imaged on a LSM 700 laser scanning confocal microscope (Zeiss), with Z-stacks of 21 slices spaced at 0.99 μm. We then used QuPath v0.2.3 to quantify the overlap of FISH and IHC signals in the maximum projection images.

#### Analysis

For the c-Fos/RNAscope experiment, we assigned neuron and transmitter identity to cells by defining section-specific cutoffs as the mode of the density function of the log-transformed distribution of RNA punctae number minus half (for *NeuN*) or one time (for *VGluT2*, *Gad1*) the standard deviation of the distribution of RNA punctae number. We defined cells as neurons or as excitatory/inhibitory when they had at least three *NeuN* or *VGluT2*/*Gad1* punctae, respectively, and exceeded the section-specific cutoffs. From this dataset, we then calculated the following three variables: percent of neurons that co-express a given transmitter, percent of transmitter-positive neurons that co-express c-Fos, and enrichment ratio (percent of transmitter-positive neurons that co-express c-Fos, divided by percent of neurons that co-express c-Fos). For the complete dataset, we then generated a mixed-effects linear model [response ~ (variable + species + stimulus + transmitter + brain region)^5^ + section ID] using the R package lme {lme4}, and evaluated the model by contrasting stimulus (percent of transmitter-positive neurons that co-express c-Fos) or species (percent of neurons that co-express a given transmitter, enrichment ratio) with emmeans {emmeans} and contrast {emmeans}. We adjusted P-values with the Benjamini-Hochberg method.

### *In vivo* Neuropixels probe recordings

#### Head-post surgery

Three animals of each species (2-4 months old) were anesthetized with isoflurane (Iso-vet; 3% for induction, 1-3% during surgery), placed into a stereotaxic system (Narishige, SR-5N), and given dura tear (Novartis, 288/28062–7) to protect their eyes. After removing the hair on the head with depilation creme, we injected lidocaine (Xylocaine 0.5%, 0.007mg/g) under the skin above the skull and then incised the scalp along the midline to reveal the skull. A metal headpost was fixed on the skull using dental cement (Superbond C&B, Prestige-dental). The animals received a single injection of buprenorphine (0.2 mg/kg I.P.) and Emdotrim in the drinking water for the next 3-5 days.

#### Experimental procedure

After at least 3 days of recovery, the animals were anesthetized briefly and a craniotomy above the SC was performed using a dental drill. Still under anesthesia, we transported the animals to the recording setup, where they were fixed with their headpost on a ball floating on air (polystyrene white ball, 20 cm diameter). In some cases, we repeated recordings again on later days; in these cases, we briefly anesthetized the animal in its cage and transported it to the recording setup.

A Neuropixels probe phase 3A (imec, Belgium (Jun *et al*. 2017)) coated with a fluorescent dye (DiD, DiI or DIO, Thermofisher) was lowered slowly into the right SC and dPAG. We targeted the center of the SC based on anterior-posterior coordinates and the midline to detect responses to the upper visual field and in the dPAG. We then covered the exposed brain and skull with artificial cerebrospinal fluid (150mM NaCl, 5mM K, 10mM D-glucose, 2mM NaH2PO4, 2.5mM CaCl2, 1mM MgCl2, 10mM HEPES adjusted to pH 7.4 with NaOH).

This setup allowed animals to walk and run on the ball, with tight control over their field of view. After 20 mins, we presented visual stimuli on a 32-inch LCD monitor (Samsung S32E590C, 1920×1080 pixel resolution, 60 Hz refresh rate, average luminance of 2.6 cd/m^2^) placed 35 cm in front of the animal’s left eye (covering 90° of azimuth and 70° of altitude) while recording the neural activity and the movement of the animal on the ball with two motion sensors (Tindie, PMW3360). We recorded the 384 electrodes (16 µm lateral spacing, 20 µm vertical spacing) at the tip of the probe, covering 3840 µm in depth. Signals were recorded at 30 kHz using the Neuropixels headstage (imec), base station (imec), and a Kintex-7 KC705 FPGA (Xilinx). High frequencies (>300 Hz) and low frequencies (<300 Hz) were acquired separately. To select the recording electrodes, adjust gain corrections, observe online recordings, and save data, we used SpikeGLX software (https://billkarsh.github.io/SpikeGLX). We simultaneously recorded the timing of visual stimulation using digital ports of the base station.

Following these recordings, the animals were euthanized, and the probe location was verified by confocal images of the fluorescent dye in 200 µm thick slices stained with DAPI. We included recordings only in cases in which the probe went through the sSC and dSC as well as the dPAG, and in which we could detect clear light responses in the SC.

#### Visual stimuli

Visual stimuli were presented on a gray background and were controlled by Octave (GNU Octave) and Psychtoolbox (http://psychtoolbox.org; (Brainard 1997; Pelli 1997)). Here, we analysed visual responses to 10 repetitions of a black looming disk (from 4° to 50° visual angle in 0.3 s; the disk stayed at full size for 0.5 s before a 3 s gray background) and a dimming disk that stayed at a size of 50° visual angle and changed from gray to black within 0.3 s. All animals were tested under dim daylight conditions (normal screen brightness or 1 log unit darker). For some animals, we conducted additional recordings under moonlight conditions (3-4 log units darker). We used all light conditions to test for a correlation between running speed and neural activity; however, we used only daylight conditions for visual response analysis. To measure contrast response curves, we displayed looming disks of different contrasts in a randomized order with randomized inter-stimulus times of 3-7 s, resulting in 8-12 repetitions of each contrast (5, 15, 25, 35, 45, 60, 75, 90, 97% Weber contrast).

#### Analysis

##### Spike sorting

We sorted the high-pass filtered neural data using Kilosort2 (Stringer *et al*. 2019) https://github.com/MouseLand/Kilosort/releases/tag/v2.0), followed by manual curation in phy2 (https://github.com/cortex-lab/phy). Units were labelled as a real unit based on their waveform shape and auto-correlogram. We used cross-correlograms to identify spikes in different units that belong to the same cell. For subsequent analysis, we used only single unit data. We identified borders between the sSC and dSC as well as the dSC and dPAG using histological sections, local field potentials, and spiking activity.

##### Contrast response curves

To calculate contrast response curves, we used looming responses of neurons in the sSC. First, we calculated firing rates during each contrast in 100 ms bins and the subtracted background activity before stimulus onset. Then, we averaged firing rates at each contrast and normalized the data by setting peak firing rates to no stimulation (0% contrast) to 0 and the maximal firing rate at any other contrast to 1. We identified responding neurons as cells with (a) at least 10 significant responses at any contrast (out of 90 total stimulations), (b) responses to the highest presented contrasts, and (c) no sudden response drop at intermediate contrasts, while responding to lower and higher contrasts.

##### Looming selectivity index

We calculated preferences for looming or dimming stimuli from full-contrast stimuli. We calculated firing rates as the number of spikes in 20 ms bins and extracted average peak firing rate (P_l_ for looming and P_d_ for dimming) during multiple repetitions of the stimuli. We defined the looming selectivity index (LSI) as: LSI = (P_l_ – P_d_)/ (P_l_ + P_d_).

##### Locomotion events

To analyse neural activity during different movement events, we binned the measured running speed to achieve the same temporal resolution as the neural activity (100 ms bins) and normalized it such that “no movement” is set to 0, maximal forward running speed set to 1, and maximal backward running speed to −1. Then, we identified time points of onset of forward running and stopping from running. We defined the onset of forward running as: an acceleration of >0.2 within 200 ms after a speed of <0.05. We defined stopping as: a deceleration (negative speed difference of >0.1) from a speed of >0.1 to a speed of <0.05. For analysis of locomotion, we included only events that were not preceded by a visual stimulus onset in the previous 1s.

##### Neural activity during locomotion

We calculated the z-score of neural activity of sorted single units in the dPAG in the 7 s before and after each event onset. The z-score was calculated as the firing rate binned in 100 ms bins minus the mean firing rate and divided by the standard deviation across the entire recording. For representative neurons, the firing rate was calculated in 20 ms bins. We calculated firing rates and z-scores around the onset of each looming stimulus in the same way.

##### Locomotion/activity correlation

To correlate movement events and the corresponding neural activity, we calculated the mean neural activity during each event (onset of forward running or stopping). We then calculated the correlation coefficient of the speed trace and the average neural activity using ‘corr’ in Matlab. We estimated the 95% confidence intervals per species as well as for a Gaussian distribution with mean 0 and the same variance using DABEST (Ho *et al*. 2019). Similarly, we estimated the cross-correlation of events and mean neural activity using the ‘xcorr’ function in Matlab.

##### Pre-running activity

We calculated residuals to extract neural activity that is not explained by the running speed. To achieve this, we normalized both the running speed and the z-scored neural activity in the 2 s before running event onset. Then, we subtracted the running speed from the neural activity to obtain residual spiking activity that does not follow the changes in running speed. Finally, we calculated the change in z-score between the 0.6 s before running onset and 5 to 1 s before running onset to quantify the increase in neural activity right before running. As described above, we estimated intervals and Gaussian distributions with mean 0 using DABEST.

### Optogenetic activation experiments

#### Virus injection and fiber implantation

To optogenetically activate dPAG neurons, we injected a viral vector into the dPAG, followed by implantation of an optic fiber. We followed the same procedure as for Head-post surgery. Following the craniotomy, we injected 50-100 nl of viral vector (AAV2/CamkII-hChR2(E123T/T159C)-p2A-EYFP-WPRE, UNC vector core, AV5456B or AVV2/CamkII-EYFP for control animals) bilaterally into the dPAG (*P. maniculatus*, lambda: +0.9 mm, midline: ±0.2 mm, depth: −2.9-3.2 mm; *P. polionotus*, lambda: +0.8 mm, midline: ±0.2 mm, depth: −2.6-2.9 mm) with a micropipette (Warner Instrument, G100-4) with an open tip of 30 µm attached to a microinjector IM-9B (Narishige). In two animals (one of each species), a modified injection protocol was used in which the virus was injected into the same location but at an angle of 30 degrees. In both cases, we lowered the micropipette to a position 0.1-0.2 mm below the targeted depth for 2 min, and then brought it up to the injection depth. After 1 min, we slowly injected the virus with a hand-wheel. After 5 min, we retracted the micropipette and closed the skin using Vetbond tissue adhesive (3M, 1469). After surgery, we provided antibiotics (Emdotrim, ecuphar, BE-V235523) via drinking water.

Either during or approximately 3 weeks after the viral injection, we anesthetized animals as described above and implanted an optic fiber (200 μm diameter, length 3.5mm, NA 0.39, Doric Lenses, B280-2304-3.5) above the injection sites (*P. maniculatus* depth: −2.5-3.1 mm, *P. polionotus* depth: −2.4-2.6 mm). Due to the large blood vessels at the midline, we first lowered the fiber into the brain lateral adjacent to the central blood vessel and then gently pushed it towards the midline and lowered it to the target depth by alternating steps of moving 100-200 µm down and up until the target depth was reached. We affixed the fiber with dental cement (Sun Medical LTD). After surgery, we injected animals with one dose of buprenorphine (0.2 mg/kg I.P.) and provided antibiotics (Emdotrim) in their drinking water for 3-5 days. We single-housed animals following surgery and gave them 7-20 days to recover before behavioural testing.

#### Experimental procedure

To test the effects of optogenetic dPAG activation on behaviour, we briefly anesthetized animals in their home cage with isoflurane, transferred them to a round arena (diameter: 43 cm) and connected them to a patch-cord with a rotatory joint (Thorlabs, RJPFL2). Animals typically woke up within 1-2 mins and started exploring the arena. We illuminated the arena with dim red light and video recorded behaviour with a camera (Point Grey Research, FMVU-03MTM-CS or Basler, acA1300-60gmNIR) positioned 53 cm above the center of the arena. We then used a DPSS laser system (Laserglow Technologies, R471003GX) or a diode laser (Laserglow Technologies, D4B2003FX) to deliver 50 Hz light pulses (10 ms on, 10 ms off) of 473 or 470 nm over 1 s through an optical fiber attached to the optic implant while the animals were freely moving in the arena. We verified laser power at the output of the patchcord without a fiber before and after recording sessions with a power meter (Thorlabs, PM100D with S130C sensor). Laser powers ranged from 0.3 to 24.1 mW.

We began experiments with low laser powers (< 1 mW output at the patch-cord) and increased laser power in steps of 1-5 mW to find a regime producing consistent behaviour. We then further investigated the effects of differing power levels below and above. Optogenetic triggers were sent manually using an Arduino when the animal was moving spontaneously through or along the walls of the arena. The session was terminated when the animals stopped cooperating (i.e., sat still for a prolonged time or started running erratically); most animals required more than one session to complete the measurements. We recorded videos at 30 Hz using pylon software (Basler) or Bonsai (Lopes *et al*. 2015). After optogenetic experiments, animals were anesthetized with isoflurane and decapitated.

#### Analysis of virus expression and fiber location

To analyse viral expression and confirm the location of the fiber, we fixed the brains of test and control animals overnight in 4% paraformaldehyde and then sliced them coronally into 200 µm thick slices with a vibratome. We washed the slices three times in PBS - 0.5% Triton and then incubated them with primary antibody chicken-anti-GFP (Thermo Fisher, A-10262 1:200) to stain for YFP that was co-expressed with Chr2 or YFP for control animals overnight at 4 °C. After washing with PBS – 0.5% Triton, we incubated slices with the secondary antibody (Alexa 488 donkey-anti-chicken, Immuno Jackson, 703-545-155) and DAPI (Roche, 10236276001) for 2h at room temperature. After washing with PBS, we mounted slices on coverslips, covered them with mounting medium (Dako, C0563), and imaged them using 10x and 63x objectives on a confocal microscope.

Using this approach, we were able to exclude animals with no or little YFP staining in the dPAG or with incorrect fiber placements (inside the dPAG or >250 µm above the dPAG). For the remaining animals, we then analysed the extent and location of the Chr2 expression and the fiber placement. First, we took confocal images of the 200 µm slice where the fiber was most visible (z-stack, 10x Objective). We then loaded raw images into Fiji (Schindelin *et al*. 2012) and identified the slice with the brightest YFP staining and with a clear fiber tract. We split the imaged channels (Chr2 and DAPI) and ran the StarDist 2D plugin (Schmidt *et al*. 2018) on the Chr2 channel (parameters: model – versatile, normalize image – yes, percentile low – 10, percentile high – 99, probability – 0.2, overlap threshold – 0.4). We measured and saved the area and X/Y position of each detected, labelled cell with an area < 1000 pixels. We loaded png versions of the maximal projections into Matlab to manually label the extent of the dPAG, the fiber tip, the outline of the ventricle and a control area without detected cells. We quantified the extent of viral infection as the number of detected cells within the dPAG below the fiber tip. Further, we quantified the area of dPAG with viral expression as % of the dPAG area with an intensity value above the mean + 2 STD control area intensity. We analysed the relationship between YFP-expression, fiber location, and running behaviour by calculating the correlation (’corr’ in Matlab) between % running trials with number of YFP-cells, distance between fiber and dPAG surface, and % of dPAG area with labelling.

#### Analysis of optogenetically induced behaviour

We extracted the head position of animals from each video with DeepLabCut (Mathis *et al*. 2018) and used this to estimate the movement speed of each animal during optogenetic stimulation. We excluded trials (i.e., laser triggers) for which the animal moved less than 10 cm/s on average during the 0.5 s before the laser trigger.

We classified each trial as “forward movement”, “slowing” or “other”. Forward movement was defined by a forward acceleration during the 1 s period of optogenetic stimulation, as determined by observing both a sharp increase in speed during the stimulus and visual inspection to ensure the movement was forwards. Slowing was defined as a continuous time period (>400 ms) with speeds below 70% of the baseline speed, confirmed by visual inspection. All remaining trials were classified as “other”. We used these classifications to calculate the percentage of forward movement and slowing trials (**Fig. 5F-G**).

In addition, we computed a Speed Index (SI) to calculate the mean speed during the main behaviour (11 video frames, 0.37 s) relative to a baseline:

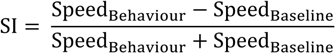

For all behaviours, the main behaviour (“Behaviour”) was centred around the maximum for forward movement trials and minimum for slowing trials. The baseline period (“Baseline”) was defined as the 0.37 s (11 frames) before the behaviour event. The SI computation resulted in positive values for forward movements and negative values for trials in which the animal slowed down.

To compare triggered and control animals as well as different laser powers, we performed two statistical tests: the two-sample Kolmogorov-Smirnov test (kstest2 in MATLAB) and estimation of effects size and confidence intervals using DABEST (Ho *et al*. 2019). Briefly, for effect size estimation, unpaired mean difference Gardner-Altman estimation was performed, in which 5000 bootstrap samples were taken and the mean difference between two groups was calculated together with the confidence interval. The p value reported is the likelihood of observing the effect size if the null hypothesis of no difference is true.

---

**Figure S1.**
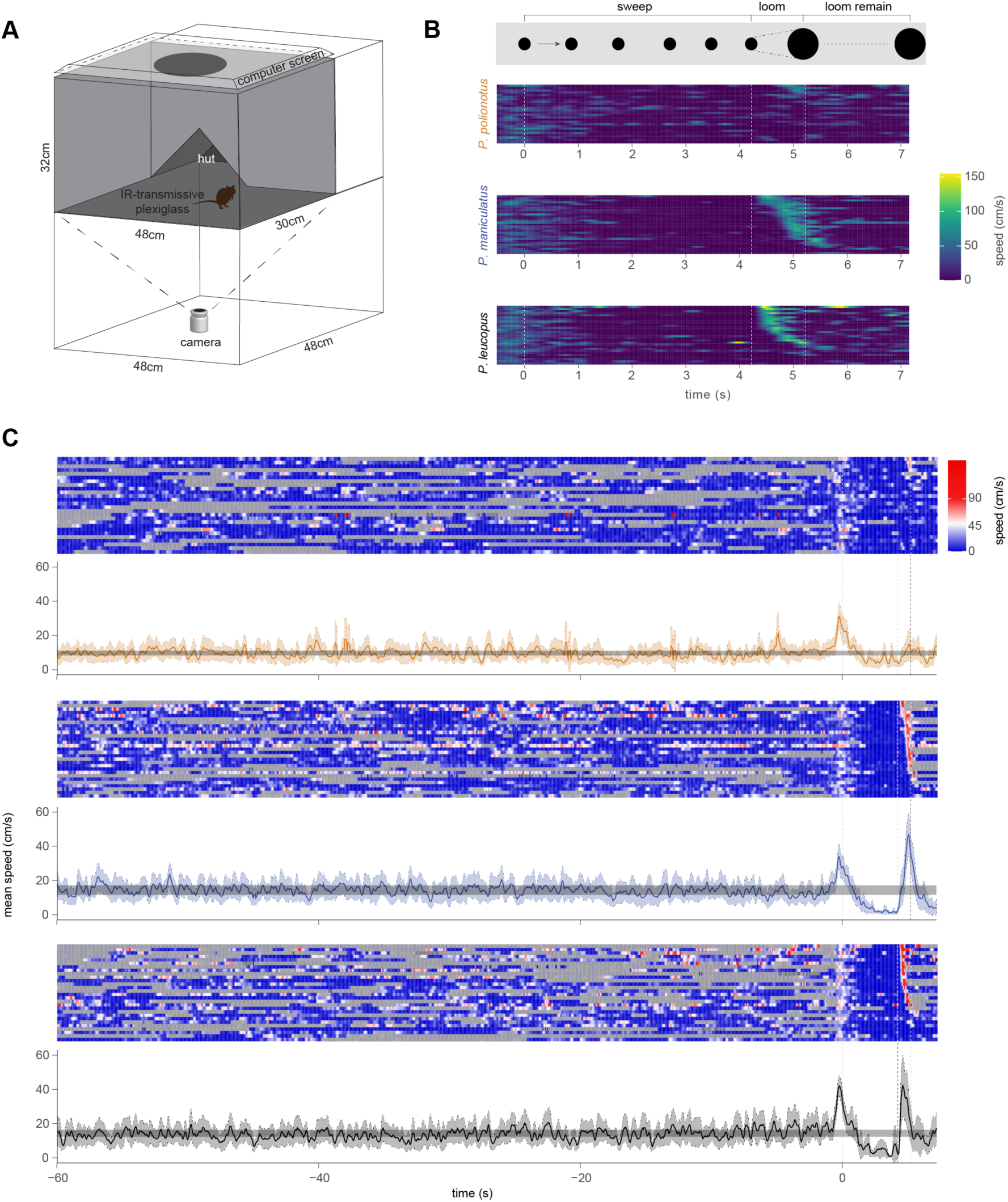
Sweep-looming experimental arena and behaviour. (**A**) Arena used to measure behavioural responses to looming stimuli in *Peromyscus* mice. Dimensions for area available to the mouse (grey) and general setup are provided. (**B**) Raster plot showing full range of mouse speed (1-150 cm/s) during the sweep-looming stimulus. (**C**) Raster plots of mouse speed during the 60 s preceding stimulus onset in addition to during the sweep-looming stimulus (top). Rows represent individual mice (*P. polionotus*, n=26; *P. maniculatus*, n= 29; *P. leucopus*, n= 28). Line plots represent mean speed (solid line) ± 95% confidence limits (color shading), and the 95% confidence interval of the mean speed averaged across the complete 60 s preceding stimulus onset is shaded (horizontal grey bar).

**Figure S2.**
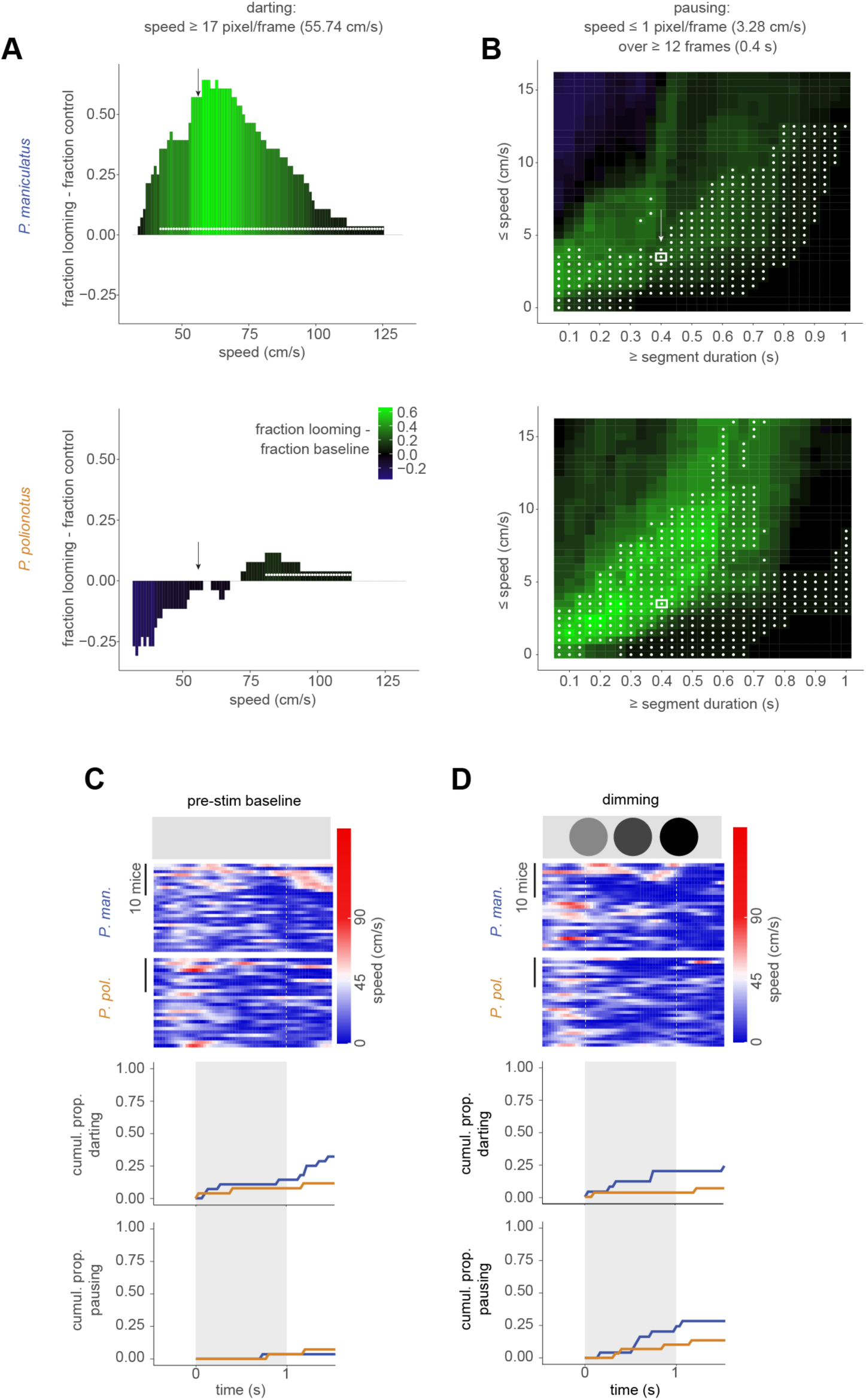
Quantitative definition of darting and pausing behaviours. (**A**) The proportion of mice that reached a given speed during looming expansion (1 s) minus the proportion of the same mice that reached the speed during the pre-stimulus control segment (1 s). A positive value indicates that more mice reached a given speed during the stimulus, and a negative value that more mice reached the speed before the stimulus. *P. maniculatus* (n=28) above; *P. polionotus* (n=26) below. White dots indicate statistically significant differences between looming-exposed and pre-stimulus proportions. Arrows indicate the quantitative threshold used to define darting. (**B**) The proportion of mice that did not exceed a given speed for a given duration while outside the hut during looming expansion (1 s), minus the equivalent proportion during the pre-stimulus control segment (1 s). *P. maniculatus* (n=28) above; *P. polionotus* (n=26) below. White dots indicate statistically significant differences between looming-exposed and pre-stimulus proportions. Arrow and white outlines indicate the quantitative threshold used to define pausing. (**C**) Raster plots (above) and cumulative proportion of darting and pausing during the pre-stimulus control segment (highlighted by vertical grey bar; below). The corresponding responses during the looming expansion are shown in Fig. 2D. (**D**) Raster plots (above) and cumulative proportion of darting and pausing during dimming stimulus (below). *P. maniculatus* (n=25); *P. polionotus* (n=30). For (**C-D**), mice are sorted by onset first of escape and then pausing, with earliest on top. Chi-Squared test; NS=not significant. Statistical significance was tested with a Bonferroni-corrected binomial test.

**Figure S3.**
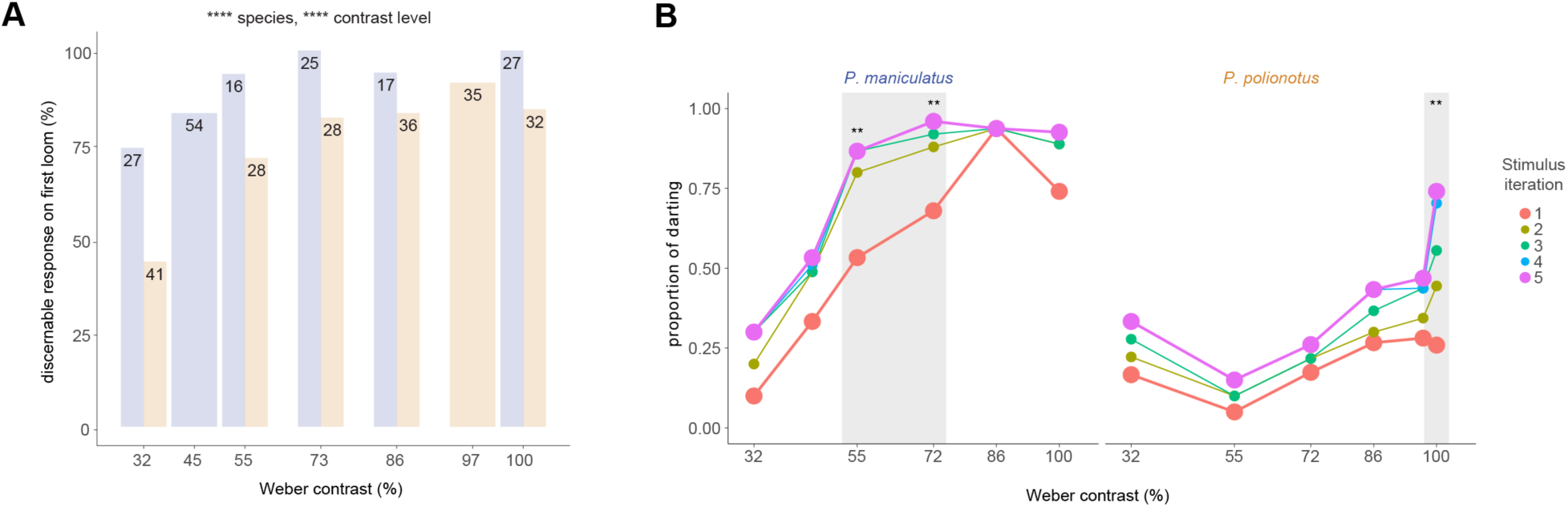
Response rate and cumulative proportion of darting by stimulus repeat. (**A**) Percentage of mice (*P. maniculatus*, blue; *P. polionotus*, gold) in each species that showed a discernable response during the first looming iteration, by contrast level of the looming stimulus (5 levels tested for both species; 2 levels tested in one species). Numbers indicate total sample size. Both contrast level and species have a significant effect on the probability of observing a discernable response. Statistical significance was tested with a logistic regression. (**B**) Of the mice that showed a discernable response during the first looming iteration, cumulative proportion of darting by stimulus iteration for *P. maniculatus* (left) and *P. polionotus* (right). The first (red) and last iteration (magenta) are highlighted with larger data points. Contrast levels with statistically significant differences among all five stimulus iterations are highlighted in grey. Statistical significance was determined by a Chi-squared test. ** *P* < 0.01; **** *P* < 0.0001.

**Figure S4.**
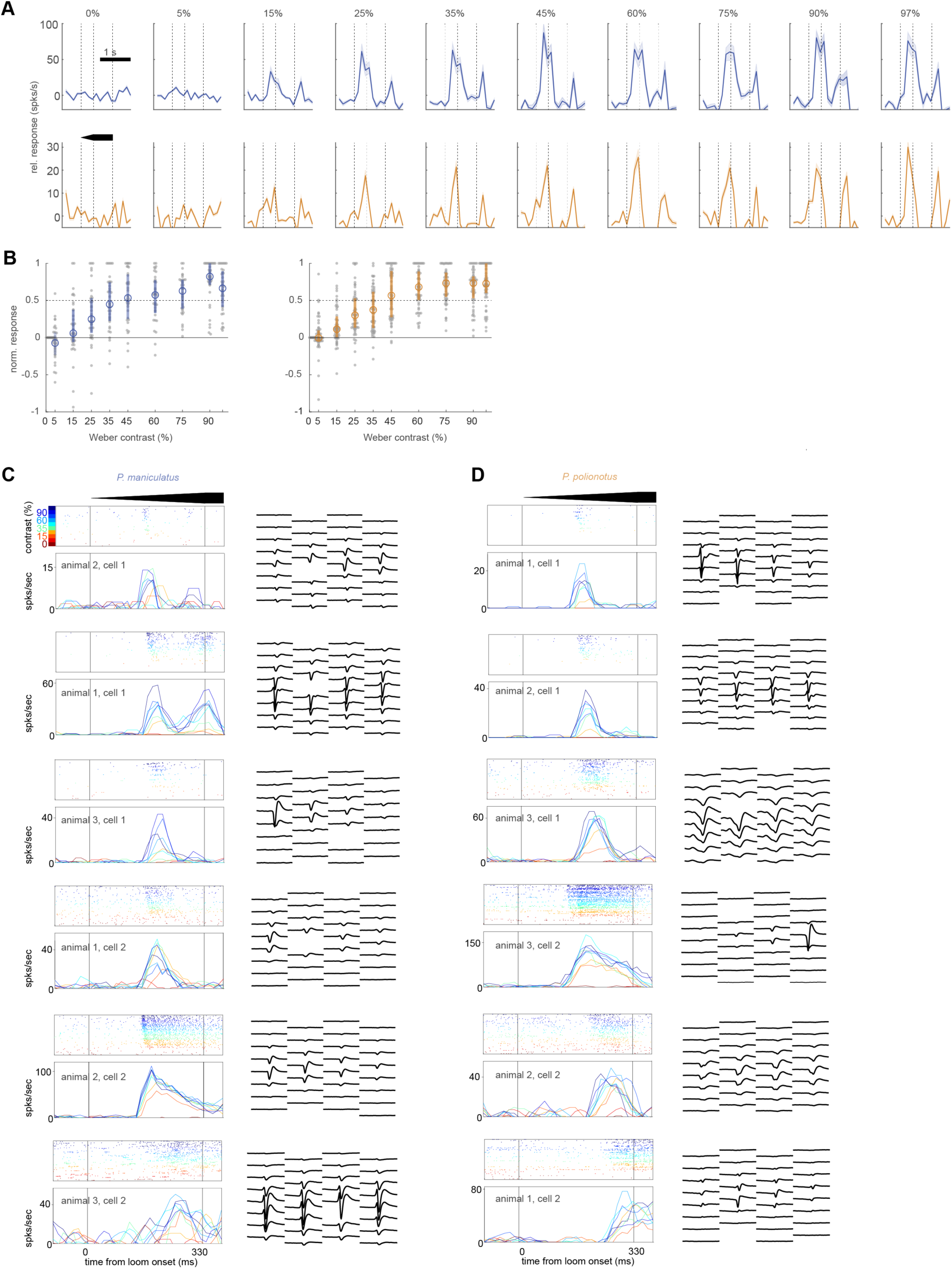
Responses of individual neurons to looming stimuli of a range of contrast levels. **(A)** Mean ± SE of responses of all sorted cells in the sSC for one *P. maniculatus* (top, blue) and one *P. polionotus* (bottom, gold) exposed to looming stimuli at different Weber contrasts (0-97%). (**B**) Normalized response strength (0 = baseline activity; 1 = maximal activity) for all responding cells in the sSC for *P. maniculatus* (blue, n=3) and *P. polionotus* (gold, n=3). Circles indicate mean, lines indicate 25-75% of data. (**C-D**) Looming responses of six example cells from the sSC of (C) *P. maniculatus* (n=3) and (D) *P. polionotus* (n=3). Raster plots and firing rates (smoothed 20 ms bins) show average response for each contrast (shown in colour code). Waveform footprint (average of 2000 waveforms per cell) on the Neuropixels probe is shown on the right.

**Figure S5.**
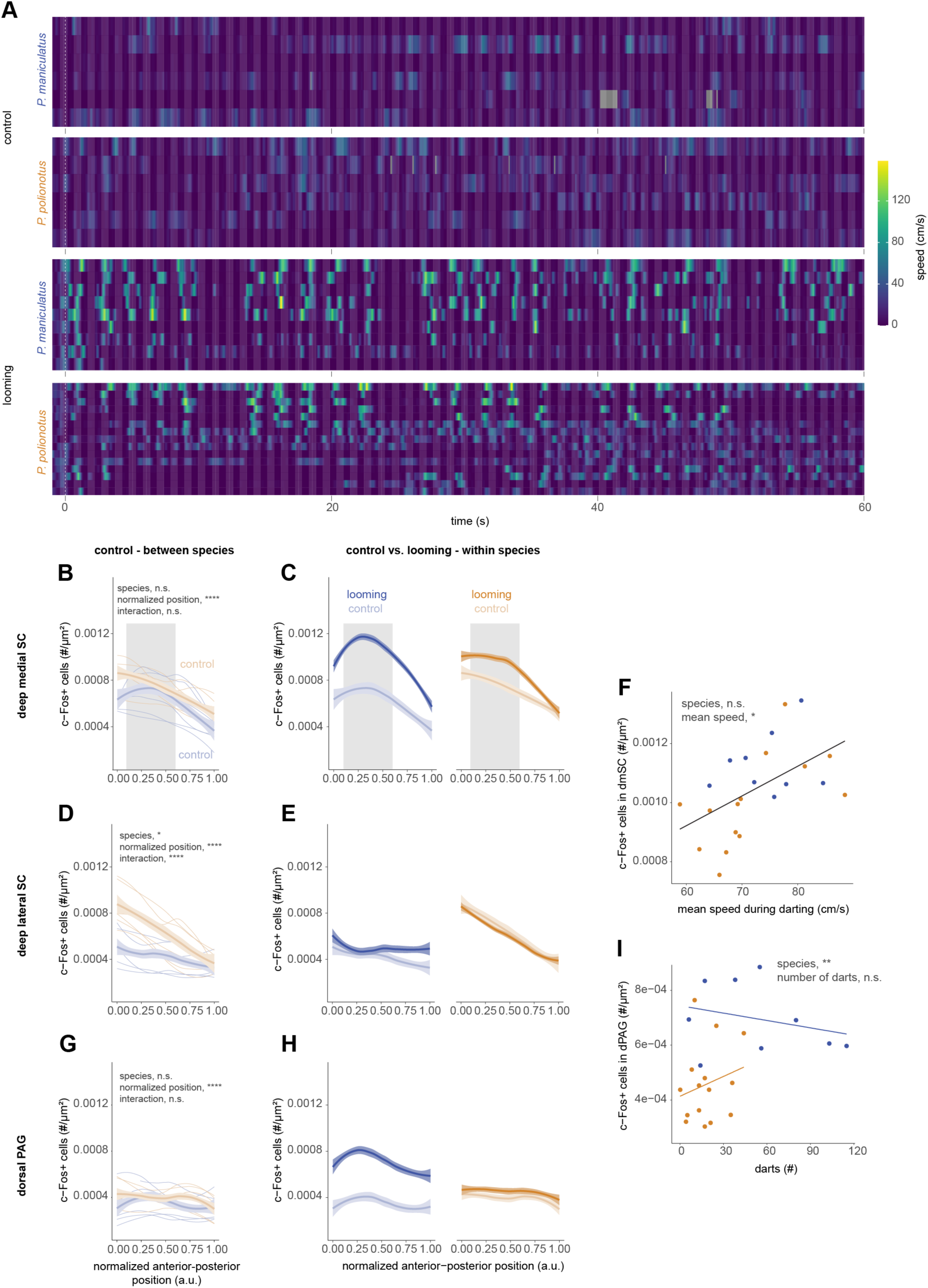
Additional information about c-Fos experiment. (**A**) Raster plot of speed of mice in control (n=6 each species) and looming (first 60 s following looming onset; *P. maniculatus*, n= 9; *P. polionotus*, n=15) assays included in the c-Fos analysis (shown in Fig. 3B). Dashed line indicates stimulus onset. (**B**) Number of c-Fos+ cells in control mice of both species (*P. maniculatus*, blue; *P. polionotus*, gold) along anterior-posterior position in dmSC. Lines represent individual mice (thin), mean per species (thick) and 95% CI (shading). Statistical significance was tested with a linear mixed effects model, including animal ID as a random effect. (**C**) Number of c-Fos+ cells in control and looming-exposed mice along anterior-posterior position of dmSC. Levels in looming-exposed mice are maximized in the central dmSC (highlighted in grey boxes). The sections within the grey boxes were used for the analyses in Fig. 3. (**D**) Same as (B), but for dlSC. (**E**). Same as (C), but for dlSC. (**F**) Number of c-Fos+ cells in the dmSC as a function of mean speed during darting (*P. maniculatus*, n=9; *P. polionotus*, n=14). Statistical significance was tested with a linear fixed effects model. (**G**) Same as (B), but for dPAG. (**H**) Same as (C), but for dPAG. (**I**) Number of c-Fos+ cells in the dPAG as a function of the number of darts. Statistical significance was determined by a linear fixed effects model. n.s. not significant; * *P* < 0.05; ** *P* < 0.01; **** *P* < 0.0001.

**Figure S6.**
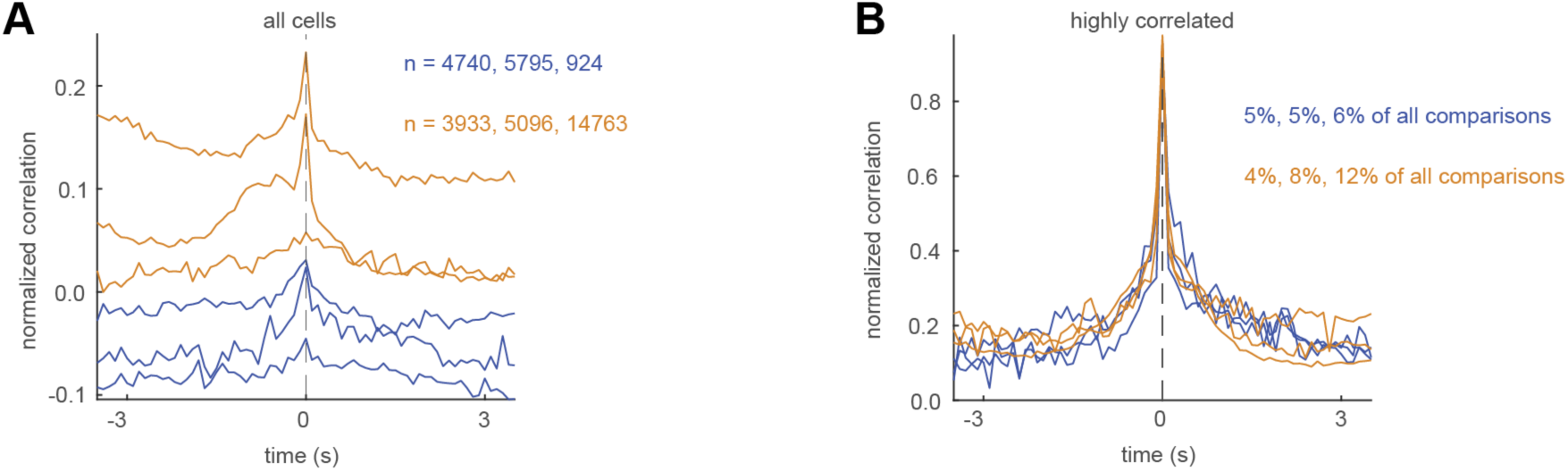
Correlation of neural activity in dSC and dPAG. (**A**) Average normalized cross-correlation of all recorded activity in the dSC and dPAG. The number of comparisons (cells in dSC vs cells in dPAG) for each animal (*P. maniculatus*, blue; *P. polionotus*, gold; n=3 for each species) is shown. (**B**) Average normalized cross-correlation of all dSC and dPAG comparisons with a correlation coefficient > 0.8 (“highly correlated”). Percentages indicate the fraction of all comparisons that fulfilled the criterion of highly correlated activity for each animal.

**Figure S7.**
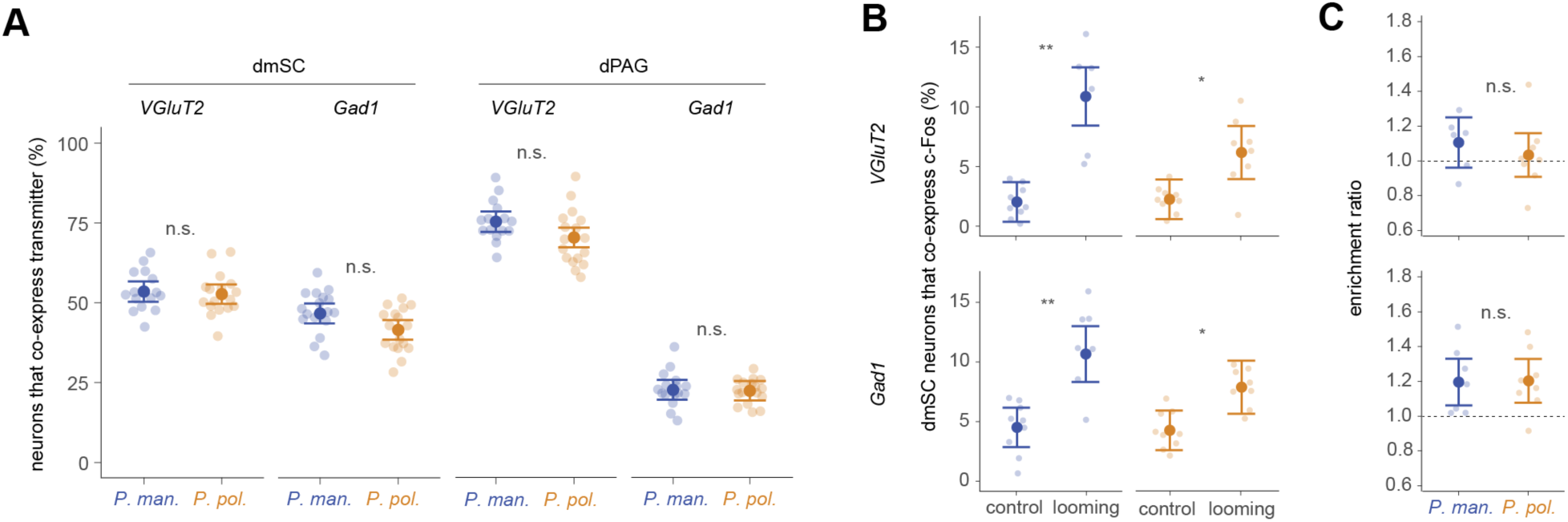
Additional information about RNAscope experiment. (**A**) Proportion of excitatory (*VGluT2+*) and inhibitory (*Gad1+*) neurons in dmSC and dPAG. (**B**) Proportion of c-Fos+ excitatory (top) and inhibitory (bottom) dmSC neurons in control and looming-exposed mice. (**C**) Enrichment index [proportion of excitatory/inhibitory neurons that co-express c-Fos, divided by the overall proportion of c-Fos+ neurons] for excitatory (top) and inhibitory (bottom) neurons in dmSC of looming-exposed mice. Statistical significance evaluated with mixed effects model. n.s. not significant; * *P* < 0.05; ** *P* < 0.01.

**Figure S8.**
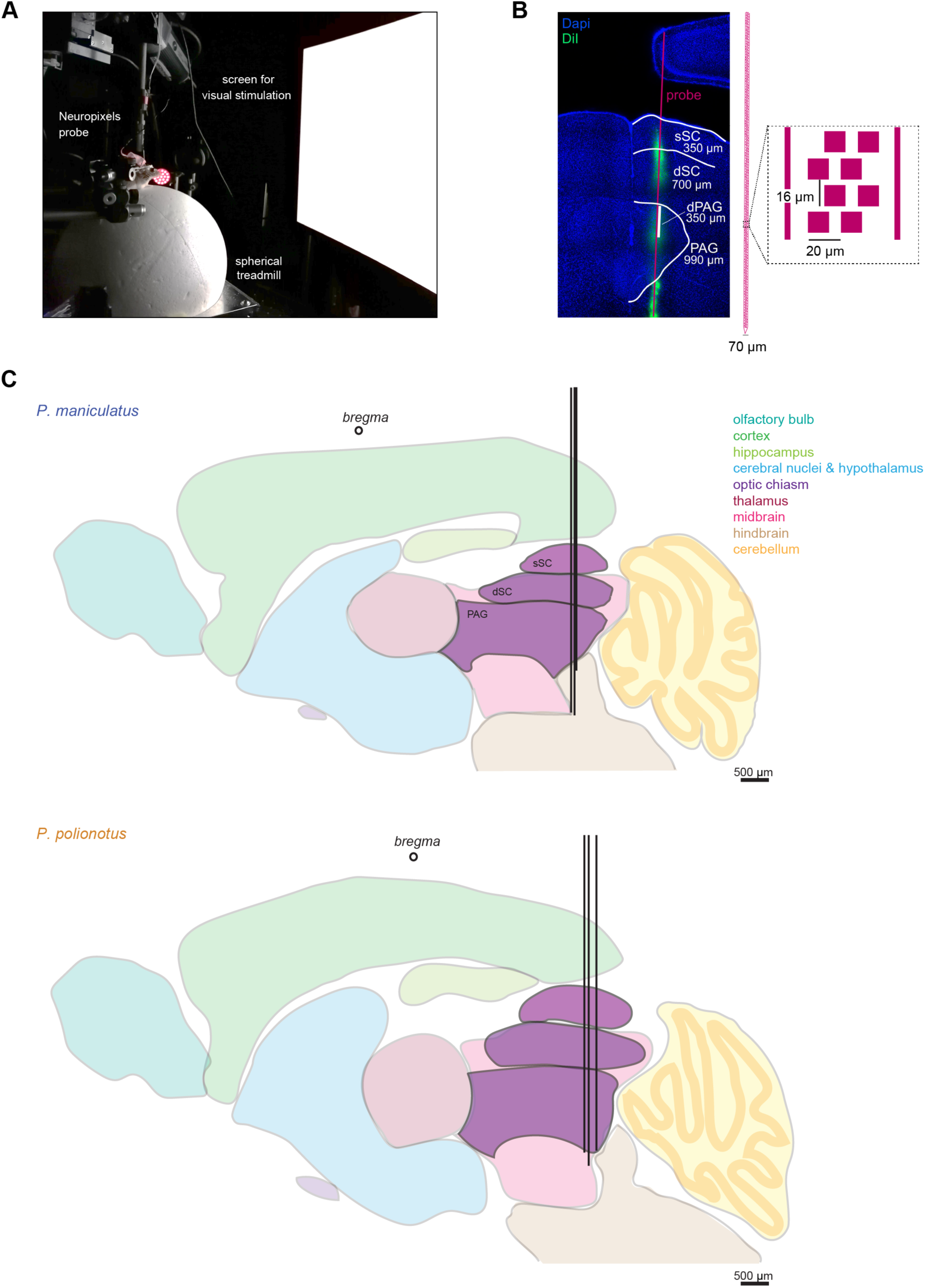
Detailed methods for the Neuropixels recordings. (**A**) Picture of a head-fixed *Peromyscus* on the spherical treadmill with implanted Neuropixels probe and computer screen. (**B**) Histological image of a coronal slice through the SC and PAG. Red line indicates the location of the Neuropixels probe that had been coated with a dye (DiI). White lines indicate separation between sSC, dSC and PAG based on inspection of the brain slice and activity patterns. Right: Bottom portion of Neuropixels probe (total: 960 electrodes) shown at the same scale as the histological image. Zoomed-in version shows positioning of individual electrodes (magenta squares). Scale bars provided. (**C**) Outlines of major brain areas in the *P. maniculatus* (top) and *P. polionotus* (bottom) brain, estimated based on *Mus* brain atlases, choleratoxin-B injections into the *Peromyscus* eye (data not shown), and multiple *Peromyscus* slices. Placement of the Neuropixels probe is indicated (N=3 of each species) as well as the anterior-posterior position of bregma.

**Figure S9.**
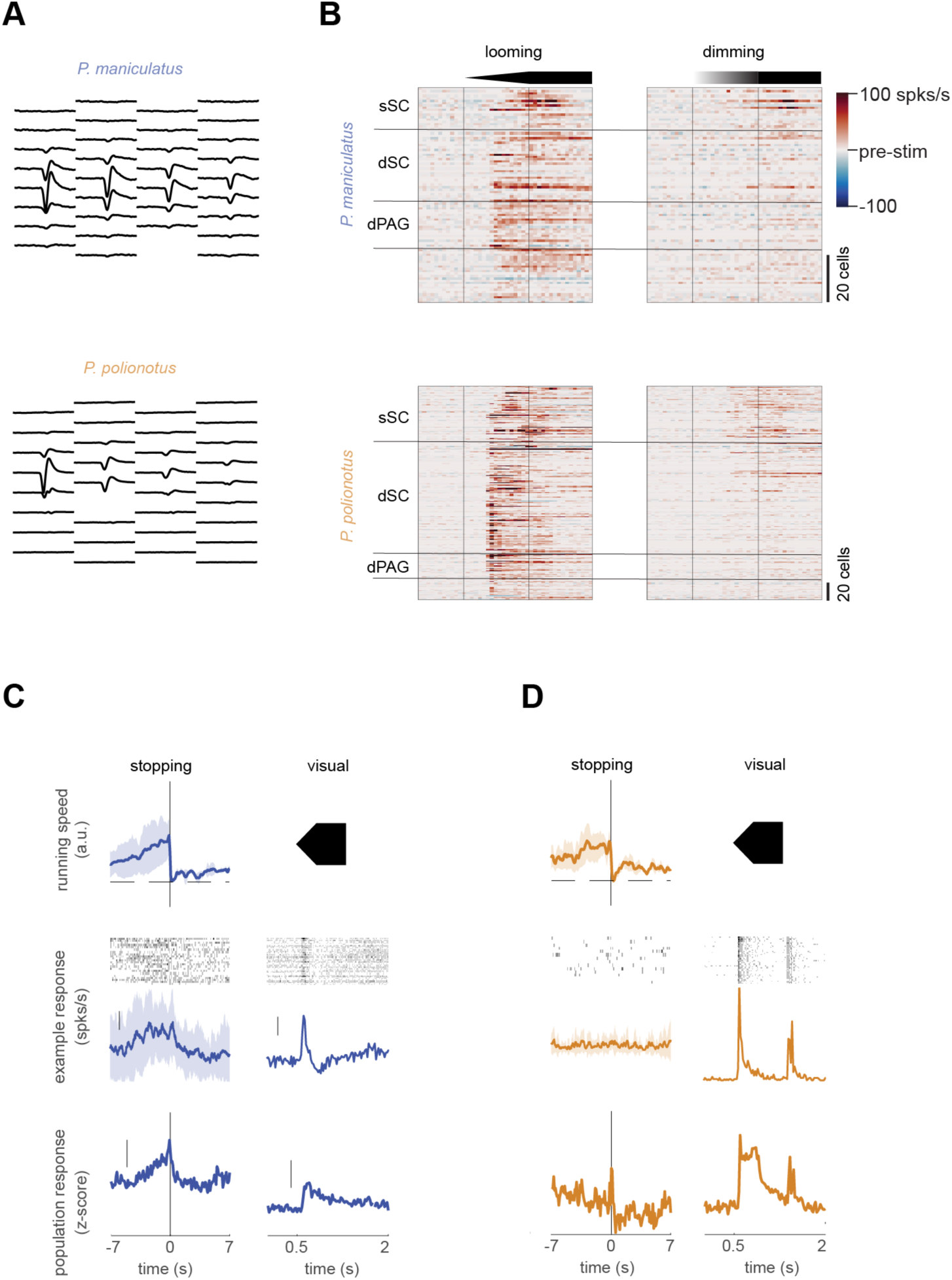
Locomotory and neural activity during stopping and visual stimulation. (**A**) Waveform footprint (average of 2000 waveforms per cell) on the Neuropixels probe of example cells from *P. maniculatus* (top) and *P. polionotus* (bottom) shown in Fig. 4B. (**B**) Average, background-subtracted looming (left) and dimming (right) response for all recorded cells shown in Figure 4C. Cells are sorted by depth; the same rows for looming and dimming correspond to the same cell. (**C-D**) Same as Figure 4D but for moments of stopping (left) and during presentation of looming stimuli (“visual”, right) for *P. maniculatus* (C) and *P. polionotus* (D). Top: Mean ± STD of the running speed during stopping events (left). Onset, expansion and duration of the looming stimulus (right). Middle: Raster plot and mean ± STD of the spiking response of a single neuron in the dPAG. Scale bar: 10 spks/s for stopping and 20 spks/s for visual stimulation. Bottom: Population response of all dPAG neurons. Scale bar: 0.1 for stopping and 0.2 for visual stimulation.

**Figure S10.**
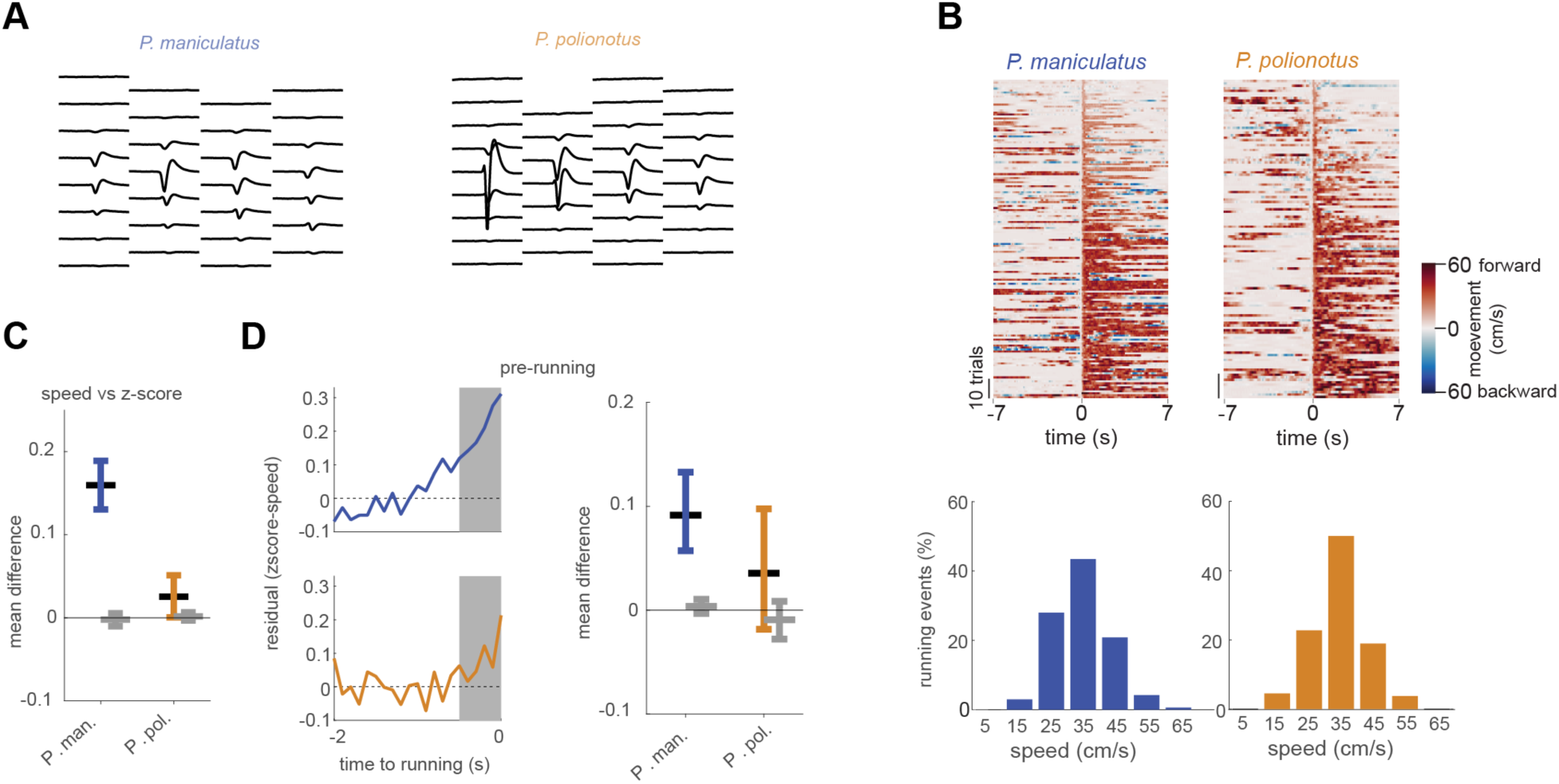
Locomotory and neural activity during running. (**A**) Waveform footprint (average of 2000 waveforms per cell) on the Neuropixels probe of example cells from *P. maniculatus* (left) and *P. polionotus* (right) taken from Fig. 4D. (**B**) Raster plot of mouse speed 7 s before and after the onset of a running event on the spherical treadmill analysed in Figure 4F, sorted by peak speed (top). Each row represents one running event. Distribution of peak speeds during running events (bottom). (**C**) Mean difference and 95% confidence intervals of the data shown in Figure 4F (in colour) and a Gaussian with mean 0 and the same variance (in grey). (**D**) Left: Residual of dPAG activity right before onset of running. Residuals were calculated as the difference of z-score and running speed. Grey area indicates the time used in Figure 4G. Right: Mean difference and 95% confidence intervals of the data in Figure 4G (in colour) and a Gaussian with mean 0 and the same variance as the data (in grey).

**Figure S11.**
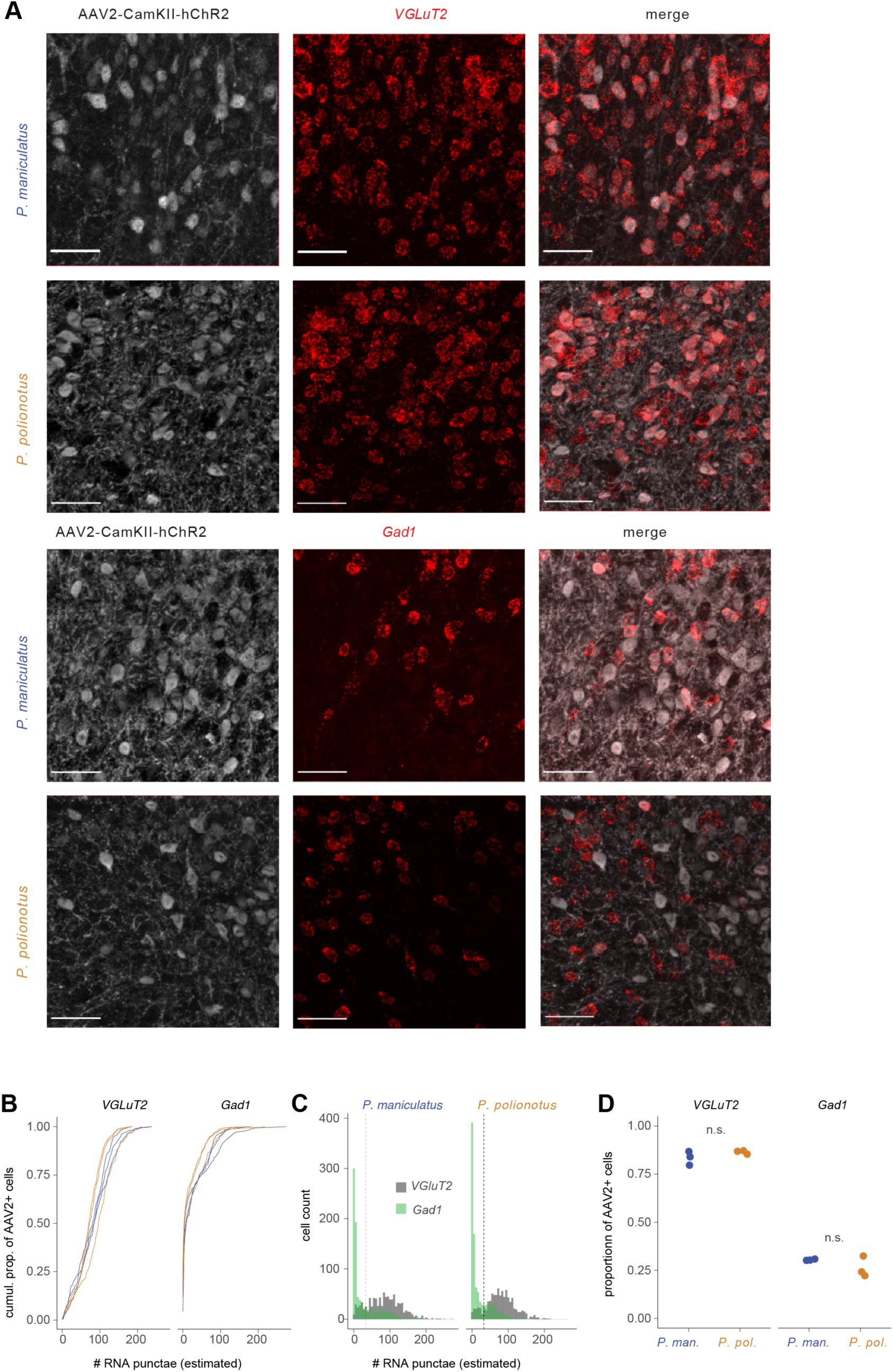
Single-molecule FISH verification of AAV infection. (**A**) Representative images of AAV2 expression and RNAscope probes against *VGluT2* (excitatory; top) and *Gad1* (inhibitory; below) for *P. maniculatus* and *P. polionotus*. Scale bar, 50 μm. (**B**) Cumulative proportion of AAV2+ cells by estimated number of RNA punctae for *VGluT2* (left) and *Gad1* (right). Individual lines represent mice (n=3, per species). (**C**) Distribution of RNA punctae across excitatory/inhibitory cells. Cut-off for assigning cell identity is indicated by the dashed line. (**D**) Percentage of AAV2+ cells that express *VGluT2* (excitatory; left) or *Gad1* (inhibitor; right) transmitter in both species. n.s. not significant.

**Figure S12.**
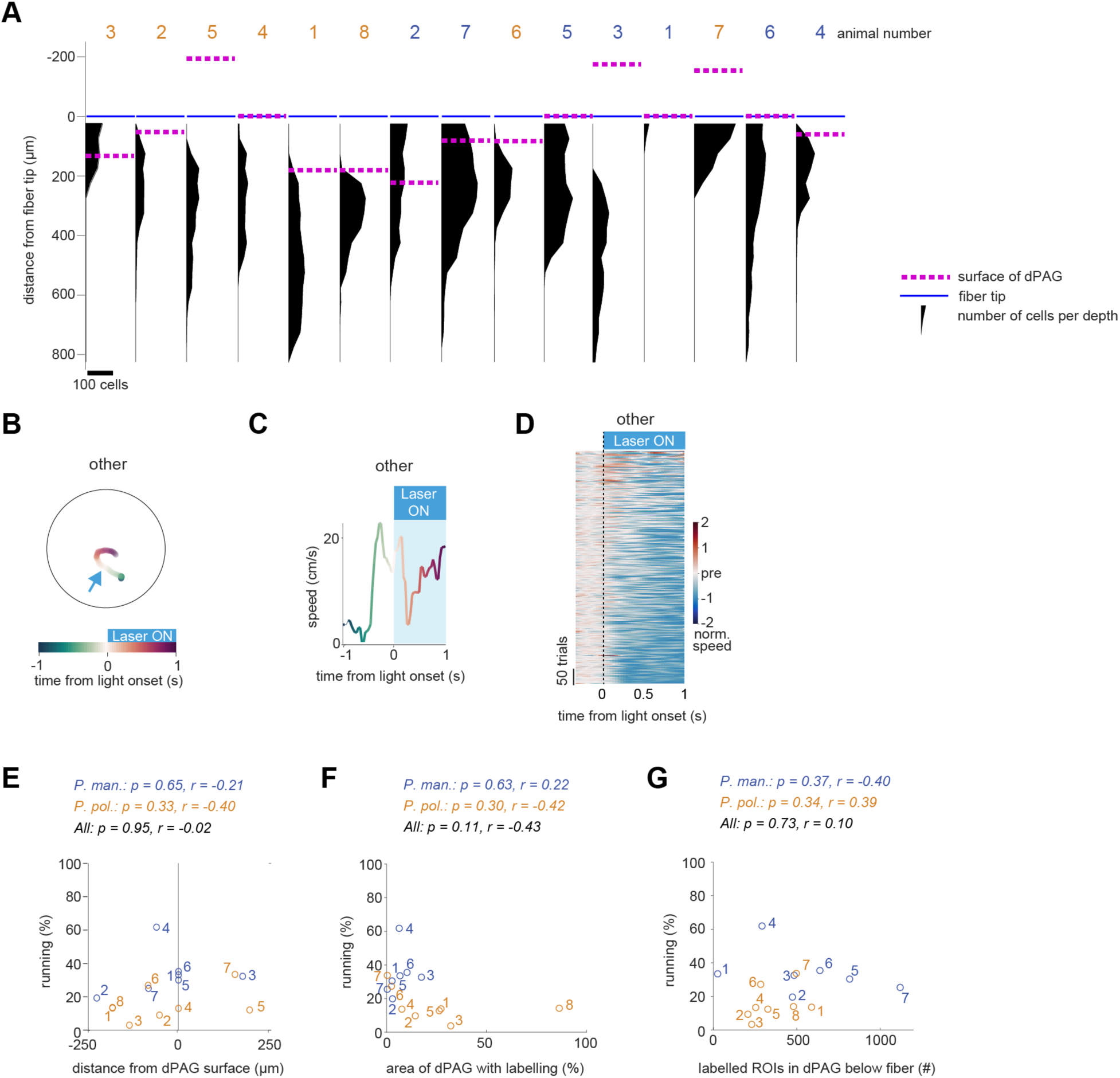
Quantification of channelrhodopsin (ChR2)-positive cells, fibre placement, and behavioural classification. (**A**) Distribution of YFP-positive regions of interest (ROIs; presumably cells) below the fiber tip. Dashed lines (magenta) indicate the surface of the dPAG; blue lines indicate the fiber tip. Animals were sorted by increasing percentage of running trials (data from Fig. 5F); animal ID numbers are provided (*P. maniculatus,* blue; *P. polionotus,* gold). Corresponding cumulative distributions of speed indices can be found in Fig. 5G. (**B**) Example trajectory of behaviour classified as “Other”. (**C**) Speed trace of example trajectory from panel B. (**D**) All traces from both species classified as “Other”. (**E-G**) Correlations between percentage of running trials and fibre location (E) and spread of the AAV virus (F-G). Correlation for *P. maniculatus* (blue), *P. polionotus* (gold), and all animals (black) is shown. Numbers indicate animal ID. (**E**) Fibre location relative to the dPAG surface and % of trials with observed running behaviour. (**F**) Percentage of fluorescently labelled dPAG area and % of trials with observed running behaviour. (**G**) Number of labelled ROIs (presumably cells) in the dPAG below the fiber and % of trials with observed running behaviour.

**Figure S13.**
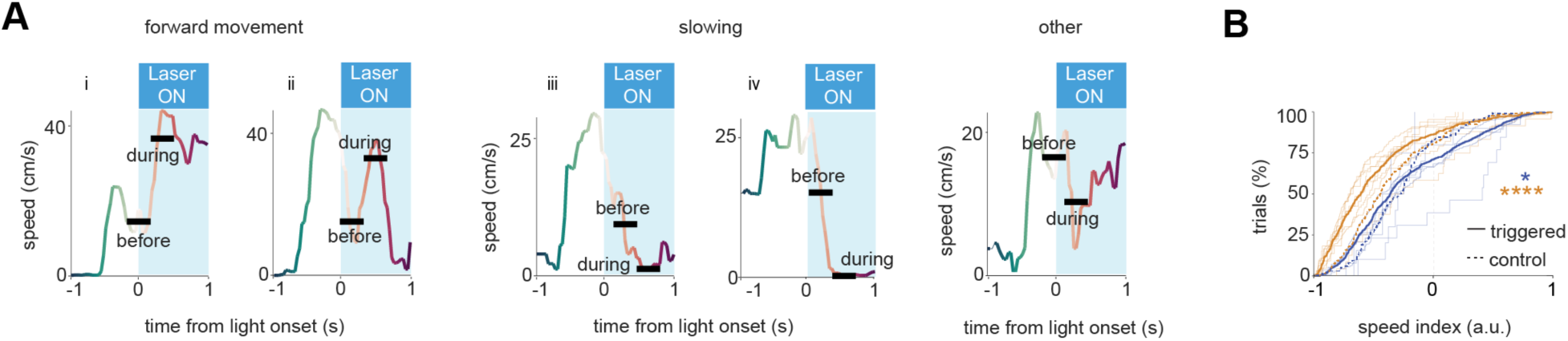
Analysis of running behaviour without classification. (**A**) Representative speed traces 1 s before and after laser stimulation for three different movement categories: forward movement, slowing, and other (following Fig. 5C). Examples of “speed before” and “speed during” intervals are shown. (**B**) Cumulative distributions of speed index of *P. maniculatus* (blue, n=7 and 706 trials total; controls: n=6, 138 trials total) and *P. polionotus* (gold, n=8, 569 trials totals; controls: n=5, 215 trials total). The speed index was calculated as ([speed before] – [speed during])/([speed before] + [speed during]), with “speed during” indicating the mean speed in the 0.37 s during the main behaviour, and “speed before” the mean speed during 0.37 s before behaviour. See Methods for details. * *P* < 0.05, **** *P* < 0.0001 two-sample Kolmogorov-Smirnov test.

